# Structural mechanism of HP1α-dependent transcriptional repression and chromatin compaction

**DOI:** 10.1101/2023.11.30.569387

**Authors:** Vladyslava Sokolova, Jacob Miratsky, Vladimir Svetlov, Michael Brenowitz, John Vant, Tyler Lewis, Kelly Dryden, Gahyun Lee, Shayan Sarkar, Evgeny Nudler, Abhishek Singharoy, Dongyan Tan

## Abstract

Heterochromatin protein 1 (HP1) plays a central role in establishing and maintaining constitutive heterochromatin. However, the mechanisms underlying HP1-nucleosome interactions and their contributions to heterochromatin functions remain elusive. In this study, we employed a multidisciplinary approach to unravel the interactions between human HP1α and nucleosomes. We have elucidated the cryo-EM structure of an HP1α dimer bound to an H2A.Z nucleosome, revealing that the HP1α dimer interfaces with nucleosomes at two distinct sites. The primary binding site is located at the N-terminus of histone H3, specifically at the trimethylated K9 (K9me3) region, while a novel secondary binding site is situated near histone H2B, close to nucleosome superhelical location 4 (SHL4). Our biochemical data further demonstrates that HP1α binding influences the dynamics of DNA on the nucleosome. It promotes DNA unwrapping near the nucleosome entry and exit sites while concurrently restricting DNA accessibility in the vicinity of SHL4. This study offers a model that explains how HP1α functions in heterochromatin maintenance and gene silencing, particularly in the context of H3K9me-dependent mechanisms. Additionally, it sheds light on the H3K9me-independent role of HP1 in responding to DNA damage.

## Introduction

Constitutive Heterochromatin is of fundamental importance to genome stability and the regulation of gene expression. It is characterized by a high copy number of tandem repeats, and an enrichment of methylation at Lysine 9 of histone H3 (H3K9me). The nuclear protein HP1, which possesses a chromodomain (CD) recognizing and binding to H3K9me, serves as a major component of the constitutive heterochromatin. Recent biophysical studies indicate that HP1 can self-associate to form phase-separated liquid droplets^1,2^, a property believed to be essential for HP1-dependent chromatin compaction. Despite these reports and extensive studies in the past few decades, how HP1 interacts with nucleosomes at the molecular level and how such interactions enable HP1 to fulfill its multifunctional role in the nucleus remain elusive. Multiple studies suggest that the CD-H3K9me interaction alone is insufficient for productive and stable HP1 interactions within the heterochromatin, and that other domains are speculated to contribute to HP1 binding to nucleosomes ^3,4^. Additional chromatin structural proteins such as histone variant H2A.Z, were also implicated in HP1 recruitment to certain heterochromatin regions in cells ^5^. Histone variant H2A.Z is predominantly found at the +1 nucleosomes at the transcription start site (TSS) of active genes, but it is also a consistent feature of pericentric heterochromatin ^6–9^. Multiple lines of studies have demonstrated that HP1⍺ directly interacts with H2A.Z-containing nucleosomes *in vivo* and H2A.Z promotes HP1⍺-mediated chromatin folding ^7,10^. Given the existing knowledge, it was proposed that H2A.Z contributes to the maintenance of the stably bound HP1⍺ population in mitotic pericentric heterochromatin ^5^.

HP1 is a highly conserved protein with three isoforms in mammals, HP1⍺, HP1β, and HP1γ. HP1⍺ and HP1β are primarily found in constitutive heterochromatin such as centromere and telomere, while HP1γ is found in both heterochromatin and euchromatin regions ^11–14^. All HP1 proteins adopt a tri-partite structure (**Fig 1A**), consisting of two globular domains, the N-terminal CD and the C-terminal chromoshadow domain (CSD). CD and CSD are conserved and homologous to each other. In addition to mediate homodimerization of HP1^15,16^, CSD has been shown to interact with proteins carrying the conserved pentapeptide motif (PxVxL)^17^. CD and CSD are connected by the variable and disordered hinge region (HR) (**Extended Data** Fig 1A). An early study showed that HR contributes to the localization of HP1 to heterochromatin^18^. Subsequent *in-vitro* studies revealed a more complex picture regarding the binding properties of HR. A cross-linking and mass spectrometry experiment of human HP1β revealed a few cross-linked sites between the HR and histone H3, although various mutations in the HR region do not significantly affect its binding to chromatin^19^. Several studies, nonetheless, confirm the ability of HR to interact with nucleic acids in a sequence-unspecific manner^18,20,21^. In additional to these three domains, the proteins have two short disordered tails at each respective end, the N-terminal extension (NTE) and the C-terminal extension (CTE) (**Fig 1A & Extended Data** Fig 1A), the functions of which remain unclear.

**Fig 1.**
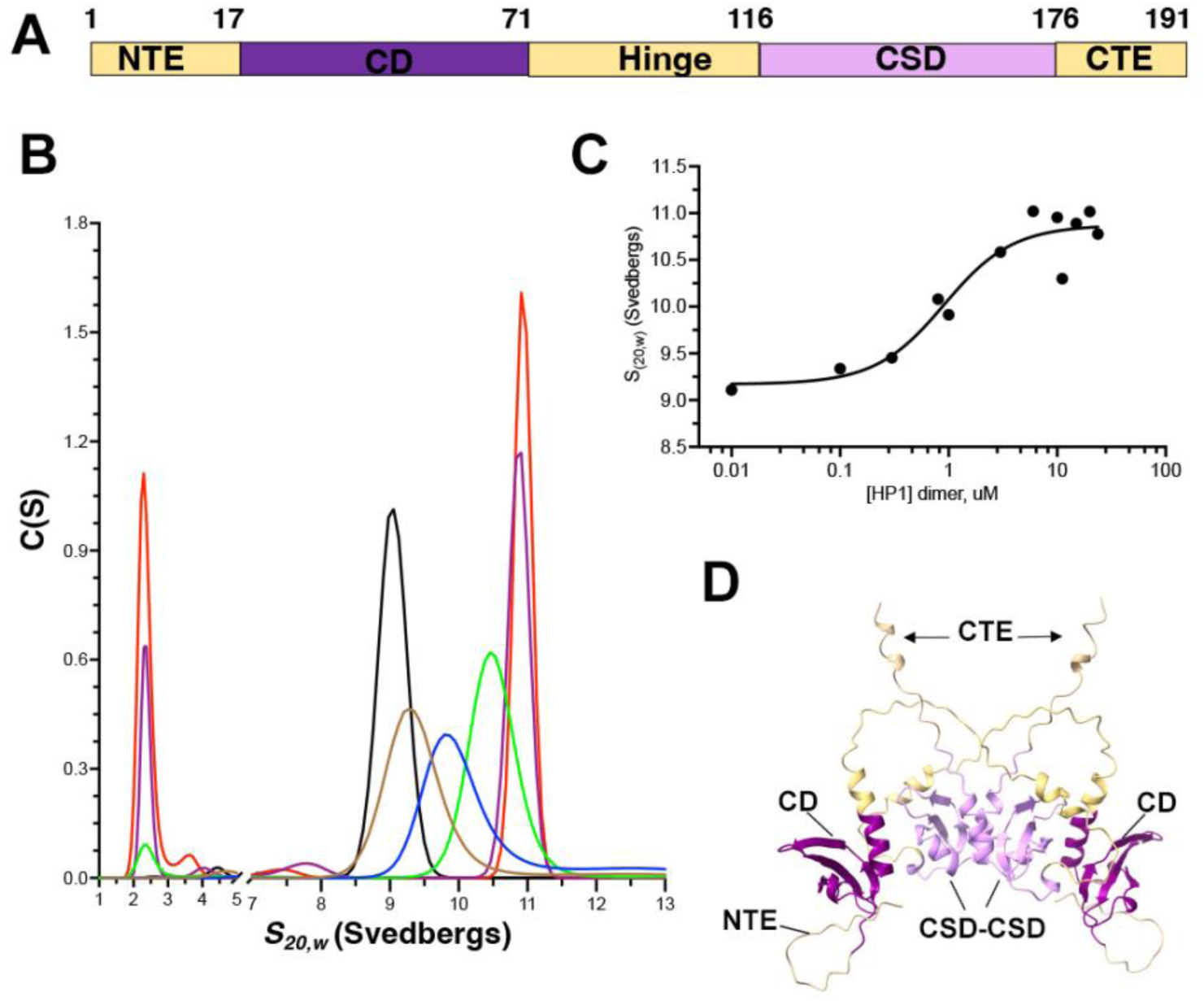
The stoichiometry of the human HP1⍺-H2A.Z nucleosome complex. (**A**) Domain architecture of the human HP1⍺ protein. **(B**) Representative c(s) distributions determined for the HP1α titration of nucleosome. The nucleosome and nucleosome – HP1α complex peaks span from 9 – 11 *S*_20,w_. The nucleosome concentration is 0.16 μM. HP1α concentrations shown are 0 (black), 0.3 (brown), 0.8 (blue), 3.0 (green), 10.0 (purple), and 20.0 (red) μM. The low *S*_20,w_ report unbound HP1α at the upper titration plateau. (**C**) The HP1⍺-nucleosome binding isotherm is well fit by the Hill equation with *R*^2^ = 0.9152, *K*_d_ = 0.85 (0.52, 1.29) μM, *n*_H_ = 1.3 (0.7, 2.5), and lower and upper limits of 9.2 (8.9, 9.4) and 10.9 (10.7, 11.4) *S*, respectively. (**D**)Alpha-fold predicted structure of human HP1⍺ dimer.

The flexible nature of the HP1 protein poses a significant challenge for structural studies of the full-length protein and of HP1 in complex with nucleosomes. Two recent cryo-EM studies revealed the low-resolution structure of HP1 in complexes with mono-nucleosomes and with di-nucleosomes^22,23^. Despite these structures, the influence of HP1-nucleosome interactions on the structure and dynamics of chromatin remains enigmatic. To gain insights into the mechanism-of-action of this important epigenetic regulator and its interplays with variant H2A.Z in heterochromatin maintenance, we applied a multidisciplinary approach to dissect the interactions between human HP1*a* and H2A.Z nucleosome. Using cryo-EM and molecular dynamic flexible fitting (MDFF) with enhanced sampling simulations we obtained a detailed model of a HP1⍺ dimer bound to an H2.Z nucleosome. The model reveals that in addition to the primary CD contacting the N-termini of histone H3 and the C-terminus of H2A.Z, the second CD interacts with the nucleosome through a new binding site near histone H2B at superhelical location 4 (SHL4). We have validated this model and confirmed both interacting interfaces with cross-linking Mass Spectrometry (XL-MS). Our model, supplemented with biochemical data, also revealed that HP1⍺ binding enhances the flexibility of terminal DNAs while simultaneously protects the internal DNA location on the nucleosome.

## Results

### The stoichiometry of the human HP1⍺-H2A.Z nucleosome complex

In the current study, we used the recombinant human HP1⍺ and H2A.Z nucleosomes containing a tri-methyl Lysine analog (MLA) at Lys9 of histone H3 to mimic the H3K9me3 modification (referred to as K9me3-H2A.Z nucleosome thereafter) (**Extended Data** Fig 1B). Previous research on the *S. pombe* counterpart, Swi6, suggested that it forms tight dimers and weak higher order oligomers in solution, with two dimers potentially binding to a single nucleosome ^23,24^.

To determine if a human HP1 protein follows a similar oligomerization pattern, we analyzed the assembly of HP1α by sedimentation velocity analytical ultracentrifugation (SV-AUC). The sedimentation coefficient and the apparent molecular weight of the dominant peak are consistent with HP1*a* being a stable dimer over the analyzed range of protein concentrations (**Extended Data** Fig 2). The median value of *M*_w_ resolved for the dominant peak of the six distributions is 42,876 Da, within 95% of the value of a HP1α dimer calculated from the protein sequence. The two minor peaks at lower and higher S_20,w_ values are consistent with HP1*a* monomer and tetramer, respectively.

The c(s) distribution for the nucleosome is dominated (92%) by a symmetric peak for which values of *S*_(20,w)_ = 9.1 and *M*_w_ = 232,578 Da is resolved. The *M*_w_ value is 99% of the value calculated for the assembled DNA and histones of the nucleosome. Only trace amounts of faster and slower peaks are evident in the distribution (**Fig 1B**). Titration of HP1α into a constant amount of nucleosome resulted in a series peaks of increasing *S*_20,w_ that plateau at *S*_20,w_ = 10.9 (**Fig 1B &C**). That single, broad intermediate peaks are observed is consistent with rapid, reversible exchange of HP1α and nucleosome. At the highest concentrations of HP1α, increasing amounts of unbound protein are visualized as a low sedimentation rate peak (**Fig 1B, orange, magenta, and green lines**). An average *M*_w_ = 329,506 Da was calculated for the dominant peak of these three highest HP1*a* concentrations at the upper plateau is 101% of the value calculated for the nucleosome complexed with two HP1α dimers (325,987 Da). Taken together, the SV-AUC results are consistent with one nucleosome binding two HP1α dimers with ∼ 1 μM affinity. The precision of the best-fit isotherm is insufficient to distinguish if HP1α binding to the nucleosome is cooperative.

### Cryo-EM Structure reveals asymmetric binding of HP1⍺ dimer on the nucleosomes

To gain structural insights of the HP1*a* –H2A.Z nucleosome complex, we conducted single-particle cryo-EM study. We employed the Grafix method ^25^ to stabilize and purify the complex. To maximize the proportion of complexes in the sample, we only collected fractions that showed clear complex formation on native gels and used them for vitrification (**Extended Data** Fig 1C). Multiple protein preparations were used for single-particle cryo-EM experiments, resulting in a substantial dataset (**Extended Data** Fig 3A **& C**). Although certain 2D class averages reveal blurry densities on both sides of the nucleosome (**Extended Data** Fig 3B), most classes in multiple rounds of 3D classification showed HP1 densities on only one side of the nucleosome (**Extended Data** Fig 3C), representing one bound HP1*a* dimer. We surmise that the apparent difference in complex stoichiometry shown in the cryo-EM data and the AUC results likely arises from the limitation of image analysis, where simultaneously aligning and resolving two flexibly bound HP1*a* dimers on the same nucleosome remains challenging.

Consensus 3D refinement of the best class produced a density map with an average resolution of 4.1Å (**map 1 in Extended Data** Fig 3C **& Extended Data** Fig 4), with the HP1*a* density ranging from 5-9 Å (**Extended Data** Fig 4A). To improve the map, we performed signal subtraction and focused refinement with two overlapping masks. One mask contains only the nucleosome, while the other contains the HP1⍺ density along with a partial nucleosome (**Extended Data** Fig 3C). The nucleosome was resolved to an average resolution of 3.9 Å (map 3 in **Extended Data** Fig 3C **and Extended Data** Fig 4). Although no improvement in overall resolution was observed (**Extended Data** Fig 4E), map 2 shows clear secondary structural features that was missing in map1.

Upon initial inspection of map 1 and map 3, we noticed that their DNA ends were notably shorter than usual (**Fig 2C**). We speculate that this feature is a result of HP1⍺ binding to nucleosomes rather than the incorporation of variant H2A.Z or the reconstitution condition. To test this hypothesis, we determined a 3.5 Å cryo-EM structure of the free K9me3-H2A.Z nucleosomes prepared under the same condition (**Extended Data** Fig 5). A comparison of the nucleosome structures revealed only about 107 bps DNA in the HP1*a* –nucleosome complex, while 130 bps of DNA were resolved in the free K9me3-H2A.Z nucleosome model (**Fig 2C**). Additionally, the HP1⍺-nucleosome complex displayed symmetrical DNA ends, while the free K9me3-H2A.Z nucleosome contains asymmetric DNA ends **(Extended Data** Fig 6). We reason that the former is the result of the binding of two copies of HP1⍺ dimers on both sides of the nucleosome in the dataset. The latter aligns with previous reports showing flexible and asymmetric terminal DNAs in nucleosomes containing histone variant H2A.Z ^26,27^.

**Fig 2.**
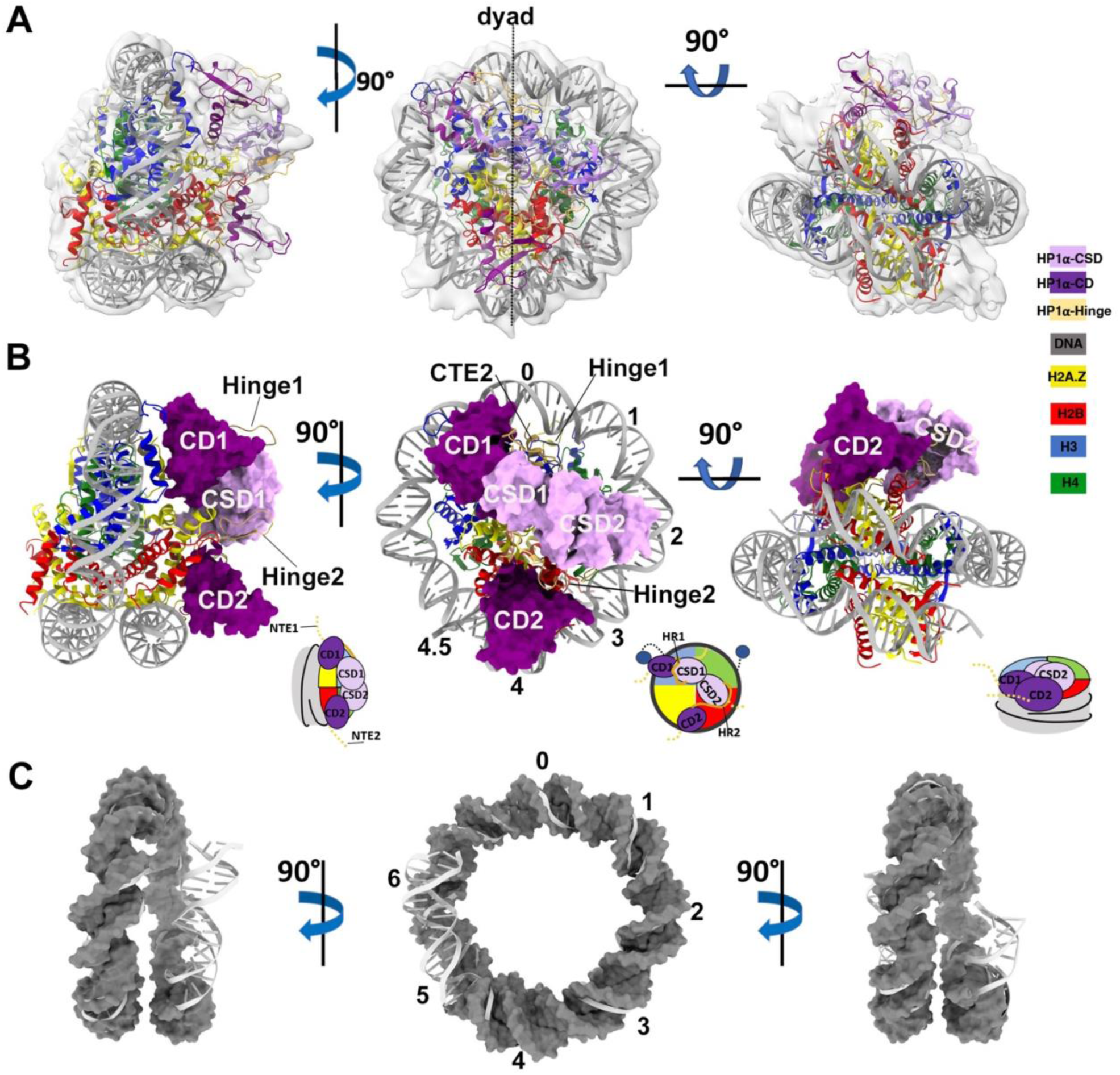
Cryo-EM Structure reveals an asymmetric HP1⍺ dimer on the H2A.Z nucleosome. (**A**) MDFF model of the HP1⍺ dimer in complex with a K9me3-H2A.Z nucleosome fitted into the consensus refined map (80% transparency),in three different views. HP1⍺ and H2A.Z nucleosome are shown in ribbon diagram. **(B**) The MDFF model color coded accordingly in the same three views as in (A). The H2A.Z nucleosome is shown in ribbon diagram and the HP1⍺ dimer in molecular surface mode. (**C**) The DNA in the HP1⍺-K9me3-H2A.Z nucleosome complex (molecular surface mode) is overlaid onto the DNA in the free K9me3-H2A.Z nucleosome (ribbon diagram). Histone cores and HP1⍺ were removed for simplicity.

To further interpret the map, we performed model building using a strategy that combine cascade or cMDFF ^28^, and Modeling Employing Limited Data (MELD) ^29^. Specifically, we used domains from the Alphafold structure of human HP1⍺ (also called CBX5) (**Fig 1D**) ^30,31^ and the crystal structures of H2A.Z nucleosome to construct initial models. The initial model was then used for model refinement and optimization in cMDFF and MELD simulation (**Extended Data** Fig 7). The resulting model revealed an asymmetric HP1⍺ dimer packed against the K9me3-H2A.Z-nucleosome (**Fig 2A & B**) through two interfaces. The primary interface was mediated by a CD (referred as CD1 in the current study) near the SHL0 position, the histone H3 ⍺N helix, and the H2A.Z C-terminal docking domain (left panel in **Fig 2B)**. The second CD of the HP1⍺ dimer (referred as CD2 in the current study) was situated on the opposite side of the dyad near SHL4 (middle panel in **Fig 2B**). CD2, along with the H2A.Z N-terminus and the H2B C-terminus in close proximity, form the second interface.

The CSD-CSD dimer in our model also covers a large nucleosome surface spanning from histone H4 near the dyad to histone H2B. However, it did not come into close contact with the nucleosome (right panel in **Fig 2B)**. It is important to note that the three disordered loops (NTE, HR, and CTE) were largely unresolved in the cryo-EM density map. Consequently, they could not be reliably modeled. Specifically, both NTEs preceding the CDs are absent from the final model, since rigid-body fitting revealed no density that could be explicitly assigned to these regions. This reflects their disordered nature despite their interactions with the nucleosome substrate. Similarly, CTE1 was not incorporated into the final model. CTE2, on the other hand, adopts an *a* helix configuration as predicted in the Alphafold structure pointing towards SHL0 (**Fig 2B**). In the model, both hinge regions are more compact (**Fig 2B**) than the predicted Alphafold structure of HP1⍺ dimer, while they largely remain disordered (**Fig 1D**).

### CSD mediates HP1⍺ dimerization without direct contact with nucleosomes

Multiple previous atomic models have shown that HP1 CSD, in isolation, form a dimer hinged on their C-terminal *a* helices with a pseudo two-fold symmetry ^32,33^ **(Extended Data** Fig 8A). Despite the limited resolution in the HP1⍺ region, map 2 shows well-resolved secondary structures in part of the CSD region (**Fig 3A-C**). This enabled us to model the CSD-CSD dimer into the density map. We applied the same CSD-CSD configuration in the crystal structures during our initial model building. The final model shows the ⍺A and ⍺B helices of each CSD monomer form a barrel-like dimer interface (**Extended Data** Fig 8A**&B**). Upon further inspection of the final 3D model, we observed notable conformational changes in the region from our initial expectations. Specifically, while CSD1 largely adopt the canonical fold of the chromo shadow domain as its *Drosophila* counterpart, structural changes were observed in the β1, β2 strands and their connecting loop in CSD2 (**Extended Data** Fig 8C). In addition, the two CSDs are positioned further apart from each other in our model when compared to the *Drosophila* HP1 CSD-CSD dimer in complex with an H3 peptide (PDB 5ti1) (**Extended Data** Fig 8 **C**) ^33^. The structural changes further accentuate the asymmetry of the CSD-CSD dimer, explaining the overall asymmetric binding of the HP1*a* dimer on the nucleosome. This observation, along with the AUC data, suggests that the HP1*a* dimeric interface in the free form is distinct from that in its nucleosome-bound form.

**Fig 3.**
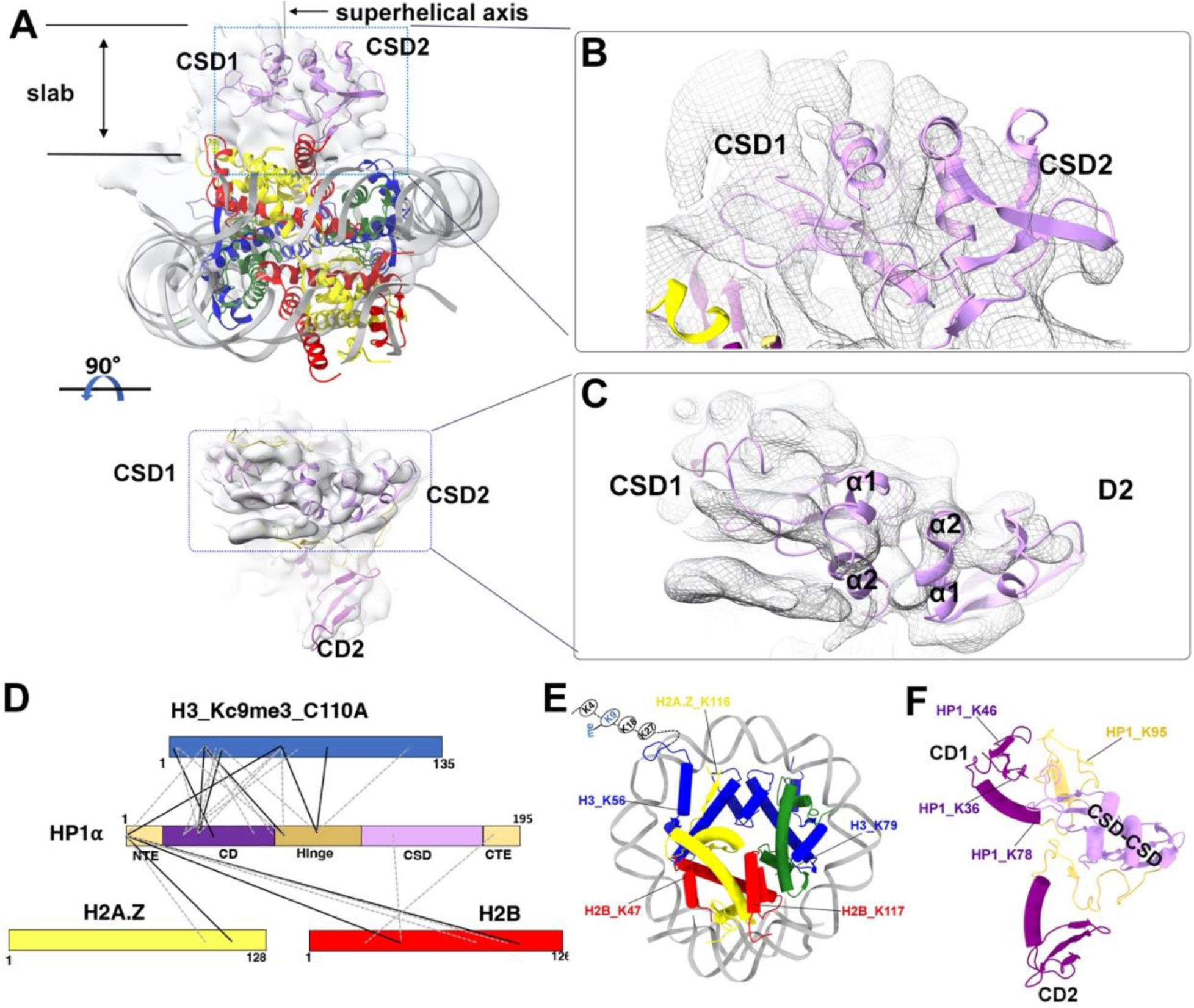
CSD mediates HP1⍺ dimerization without direct contacts with the nucleosome. **a**) The MDFF model fitted into the focused refined density map of HP1⍺-H2A.Z nucleosome (map2 in Extended Data Fig 3C) in two different views. For simplicity, the two CDs are not shown. For better clarity of the HP1⍺ dimer, only a slab of the volume in the top panel is shown in the bottom panel. The two CSD protomers are indicated and highlighted in blue dotted boxes. The map was low-pass filtered to 6 Å. The superhelical axis is indicated. **b**) Close-up view of the CSD-CSD region highlighted in the top panel in (A). **c**) Close-up view of the CSD-CSD region highlighted in the bottom panel in (A). **d**) Cross-links between HP1⍺ and histones identified by XL-MS. Solid lines are common cross-links identified from two experiments. Dotted lines are unique cross-links identified in only one of the experiments. **e**) Cross-links mapped on the nucleosome from the current model. **f**) Cross-links mapped on the HP1⍺ dimer from the current model.

The CTE, which immediately follows the CSD in HP1, has previously been found to interact with various chromatin factors ^34^. Although the precise function and the nature of such interactions remains unknown. As the exact configuration of the CTEs could not be precisely defined in the current structure, we reason that the CTE1 in the model largely remains disordered in the HP1-nucleosome complex. To validate the model and to gain further insights into the HP1-nucleosome interacting interfaces, we conducted cross-linking Mass Spectrometry (XL-MS) experiments to probe domain interactions in both free HP1⍺ and HP1⍺-nucleosome complex. In our analysis of free HP1⍺, we observed numerous inter-HP1⍺ cross-links, particularly in the hinge, CSD, and CTE regions, suggesting a propensity for self-association of HP1⍺ in these regions (**Extended Data** Fig 2C). Overall, our result implies a compact, auto-inhibited state adopted by HP1⍺ in solution, in line with previous findings ^1^,

The XL-MS experiments also revealed common cross-links between HP1⍺ and the histones. The majority of these crosslinks involve interactions between HP1⍺ and the H3 N-terminus (**Fig 3D-F**). Additionally, the N-terminal tail of HP1⍺ cross-linked with both H2B and H2A.Z C-terminus, suggesting their spatial proximity. Conversely, only a few unique cross-links, but no common cross-links, were detected between histones and CSD and CTE, indicating that both regions were further from the nucleosome surface and do not engage in stable direct interactions. Based on the structure and the XL-MS data, we further speculate that the CTEs remain mobile and accessible in the HP1⍺-H2A.Z nucleosome complex. To test this hypothesis, we conducted binding assays using a truncated HP1⍺ (HP1⍺ ΔC) from which the CTE was removed. Through electro-mobility shift assays (EMSA), we demonstrated that HP1⍺ ΔC binds to H2A.Z nucleosomes with an affinity similar to that of the full-length HP1⍺ (**Extended Data** Fig 1D), confirming our hypothesis. In summary, we conclude that the CSD contributes to the protein dimerization and HP1⍺ stability on nucleosomes without directly engaging the substrate. Our data also indicates that while both CSD and CTE can form transient interactions with the nucleosome, they do not significantly contribute to nucleosome binding.

### The interaction between CD1-H3-H2A.Z forms the primary interface between the HP1⍺ dimer and the nucleosome

Our model revealed two distinct interfaces of HP1⍺ on the nucleosome, both facilitated by CDs. The primary interface, located near the nucleosome SHL6 position, comprises CD1, histone H3 and, H2A.Z (**Fig 4A&B**). XL-MS data confirmed the existence of this interface, showing cross-links between HP1⍺ CD and H3 N-terminus, and the H2A.Z C-terminal extension (**Fig 3D)**. This interface plays a pivotal role in HP1⍺ recognizing and binding the tri-methylation at Lysine 9 of H3, as depicted in our structure. Although the CD1-H3K9me3 motif remains unresolved in our cryo-EM map, our model captures, for the first time, the close contact of the CD of a full-length HP1 dimer with histone H3 in the nucleosome. XL-MS experiments unveil cross-links not only in the flexible loop of CD1 but also in its ⍺1 helix, along with a lysine situated proximally to the hinge region (**Fig 4D**). Common cross-links on histone H3 were detected on multiple lysine at the N-terminal tail as well as on the ⍺N helix of H3 (**Fig 4D**). The close spatial proximity of these cross-linked residues, as depicted in the structural model, aligns excellently with the CD1-nucleosome arrangement in our structural model. In addition to H3, Lysine 116 of H2A.Z on its C-terminal tail is cross-linked with the HP1⍺ NTE1, as demonstrated in the XL-MS results (**Fig 3D & 4E**). This outcome not only affirms the close contact between the H2A.Z C-terminal tail and the HP1⍺ NTE1, but also suggests that the characteristic residues of H2A.Z at its C-terminus contribute to its interactions with HP1⍺ in vivo.

**Fig 4.**
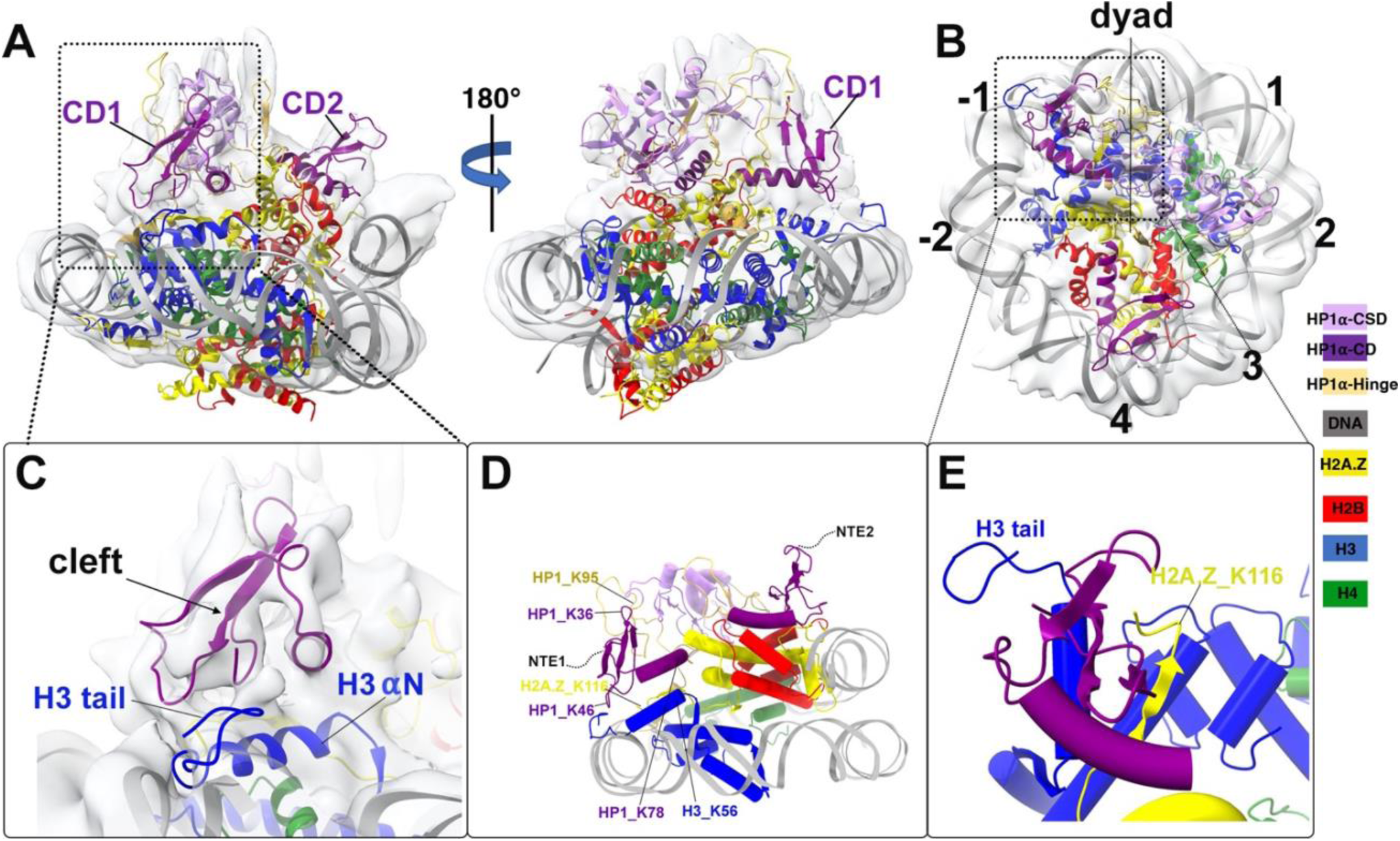
The interaction between CD1-H3-H2A.Z forms the primary interface between the HP1⍺ dimer and the nucleosome. (**A**) Ribbon diagram of the MDFF model fits into the focused refined map of HP1⍺ dimer with partial H2A.Z nucleosome (map2 in the Extended Data Fig 3 &4) in two different views. The density map was low-pass filtered to 6 Å. The primary interacting interface is highlighted in black dotted box. **(B**) The model in map 2 looking down from the superhelical axis. The SHL positions are labeled. **(C**) Close-up view of the primary interface shown in the black box in (A), showing the fit of the CD1 in the density map. **(D**) Cross-links at the primary interface identified by XL-MS. **(E**) Close-up view of the primary interface shown in the black dotted box in (B), to show the common cross-link between HP1*a* NTE1 and the H2A.Z C-terminal tail (K116).

Histone H3 ⍺N helix is known to make important contacts with the last turn of DNA at the edge of the nucleosome ^35^. Alterations of the H3 N-terminal tail and the adjoining H3 ⍺N helix have been shown to perturb DNA wrapping and histone dimer exchange on nucleosome ^36^. Furthermore, this H3-dependent nucleosome dynamics is sensitive to changes in the H2A C-terminal extension, which contacts H3 ⍺N helix to influences terminal DNA dynamics ^26,37^. Our current model demonstrated that these structural motifs constitute the primary binding site for HP1⍺ dimer. We propose that CD1 binding to this site interfere with the histone-DNA interactions mediated by H3 N-terminus and H2A.Z C-terminus, thereby increasing DNA flexibility at the edge of the nucleosome.

Additionally, we compared the CD1 in the current model to the crystal structure of its *Drosophila* counterpart (PDB 1KNE) ^38^. The comparison revealed that both structures exhibit the canonical fold of the conserved chromo domain (**Extended Data** Fig 8D). Both the *Drosophila* HP1 CD and its mouse counterpart for HP1β employ an induce-fit cleft to interact with the H3 peptide in a β sandwich conformation ^38,39^ (**Extended Data** Fig 8D). The cleft is in close proximity to the H3 tail and H3 ⍺N helix in the current model (**Fig 4C),** suggesting that HP1⍺ CD1 employs the same mechanism as other HP1 homologs in recognizing the H3K9me modification on nucleosome.

### CD2-H2B form the second interacting interface between HP1⍺ and the nucleosome

The secondary HP1⍺-nucleosome interface, composed of CD2 and histone H2B (**Fig 5A &B**), is located at a distance from the primary binding site, situated on the opposite end of the dyad and near SHL4 in our model (**Fig 2A&B**). In our model, the ⍺ helix of CD2 closely interact with ⍺1 helix of H2B, while the loop contacts the H2B ⍺C helix (**Fig 5B**). Similar to the CD1-H3 interface, the NTE2 preceding CD2 is absent in the model. According to our model, NTE2 is outward-facing (indicated as dotted line in **Fig 5C**), suggesting that it remains mobile within the complex. In our comparative analysis, we found that the CD2 in our current model closely resembles the crystal structure of *Drosophila* CD. We therefore conclude that both CDs in the HP1⍺ dimer adopts a conserved and nearly identical fold as its *Drosophila* counterpart (**Extended Data** Fig 8D).

**Fig 5.**
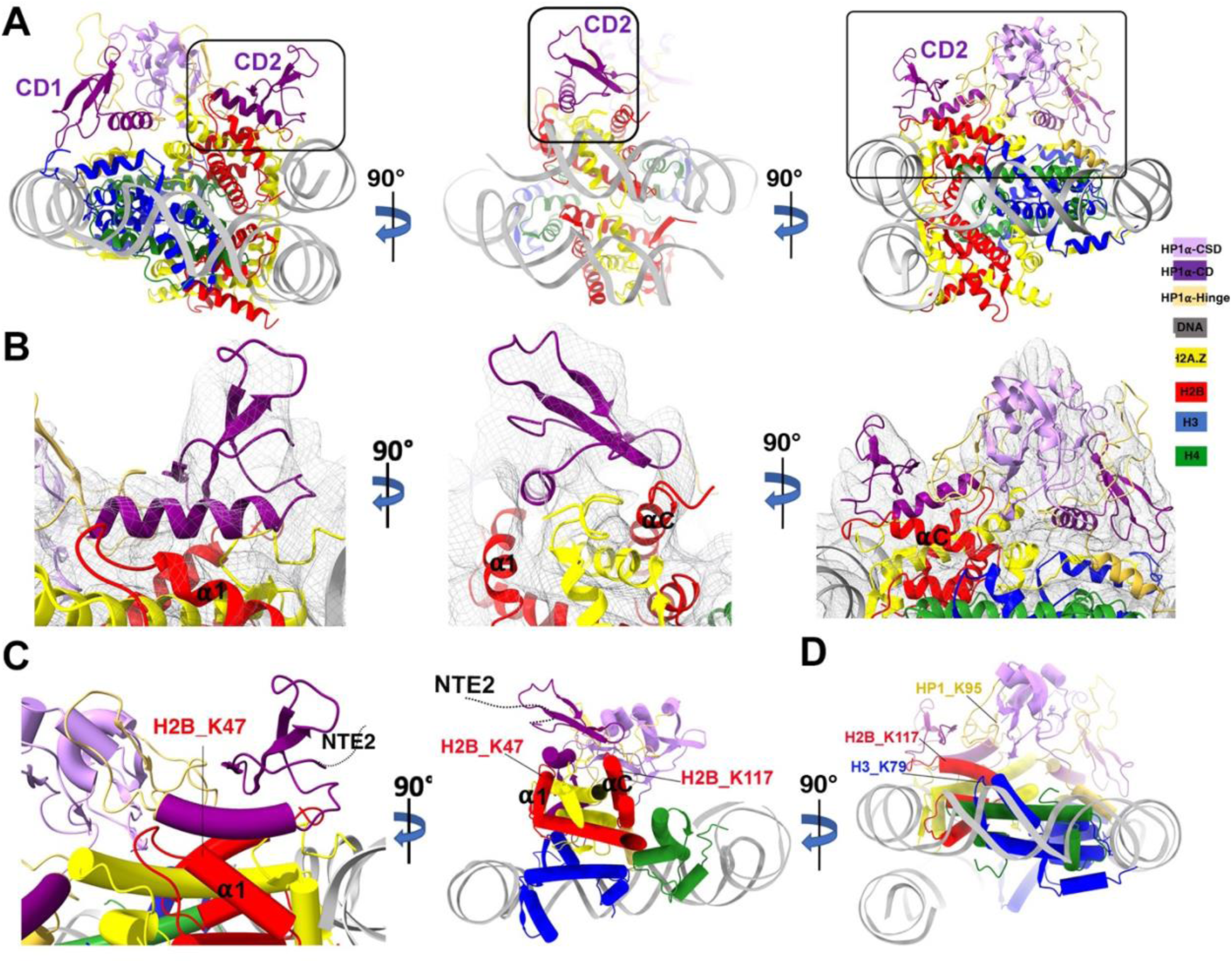
CD2-H2B form the second interacting interface between HP1⍺ and the nucleosome. (**A**) The MDFF model in three different views. The second interface mediated by CD2 and H2B is highlighted with black box. **(B**) Close-up views of the second interface as shown in (A) in its respective angle. The MDFF model was fitted into map2 (low-pass filtered to 6 Å) to show the overall fit of CD2 in the density map. **(C**) The cross-linked H2B lysine identified by XL-MS are mapped and labeled in the model (rod diagram). The views correspond to the left and middle panels in (B). The two lysine are in H2B *a* 1 helix and *a* C helix respectively. The putative position of HP1*a* NTE2 (not resolved in the model) is indicated as a dotted curve. **(D**) A cross-link between H3 K79 and a lysine at the HP1*a* hinge region (K95) is mapped in the model. The view correspond to the right panel in (B).

Our XL-MS results reveal two common cross-links and several unique cross-links between H2B and the NTE2 of HP1⍺, providing direct evidence for the existence of this interacting interface (**Fig 3D**). When mapped on the model, these common cross-links appear on H2B ⍺1 helix (K47 in **Fig 5C**) and the ⍺C helix of H2B (K117 in **Fig 5C**). This XL-MS result further validates our model. Consequently, we conclude that the CD2-H2B interactions in our model represent a novel binding interface between HP1⍺ and nucleosome. This binding site, composed of the H2B ⍺1 and ⍺C helices, is distinct from the primary binding site on H3 tail and H2A C-terminus, and it is independent of H3K9 methylations.

It is worth noting that in the current structure, the 601 Widom DNA was found to be mobile at both terminus and thus was resolved only up to SHL4.5/-4.5 location (**Fig 2C**). DNA segments near SHL4 and SHL5 are known to interact with the C-terminus of H2B and N-terminus of H2A in the major-type nucleosome ^35^. It is thus tempting to speculate that CD2 binding to these histone motifs may stabilize the histone-DNA interaction and thus protects DNA near SHL4 to SHL5.

### The effects of HP1⍺-binding on nucleosome DNA accessibility

Terminal DNA breathing is an inherent property of nucleosome dynamics. Increased terminal DNA unwrapping has been observed for nucleosomes containing histone variants H2A.Z and H2A.B ^26,27,40^. In certain structures, nucleosome complexes with chromatin remodelers and RNA polymerases have shown DNA-end deformation or peeling ^41^. To the best of our knowledge, there have been no reports of enhanced DNA breathing associated with epigenetic repressors and heterochromatin proteins. To delve deeper into HP1⍺’s potential to influence terminal DNA flexibility, as suggested by our cryo-EM model, we conducted the DNA accessibility assay using a Micrococcal nuclease (MNase). MNase cleaves DNA in a sequence-unspecific manner, allowing us to access changes in nucleosomal DNA accessibility resulting from HP1⍺ binding. Our MNase digestion of the K9me3-H2A.Z nucleosome generated a ladder-like pattern that evolved over time and eventually dissolved completely after 40 minutes (**Fig 6A**). As expected, the presence of stoichiometric histone H1, known for its role in stabilizing higher-order chromatin structures through interactions with linker DNAs, protected linker DNA and slowed down MNase digestion of K9me3-H2A.Z nucleosomes ^42–44^. This protection is evident from the appearance of two higher molecular weight bands at 200 bp and 160 bp (see **Fig 6B & C**), which were entirely absent in reactions with free K9me3-H2A.Z nucleosomes.

**Fig 6.**
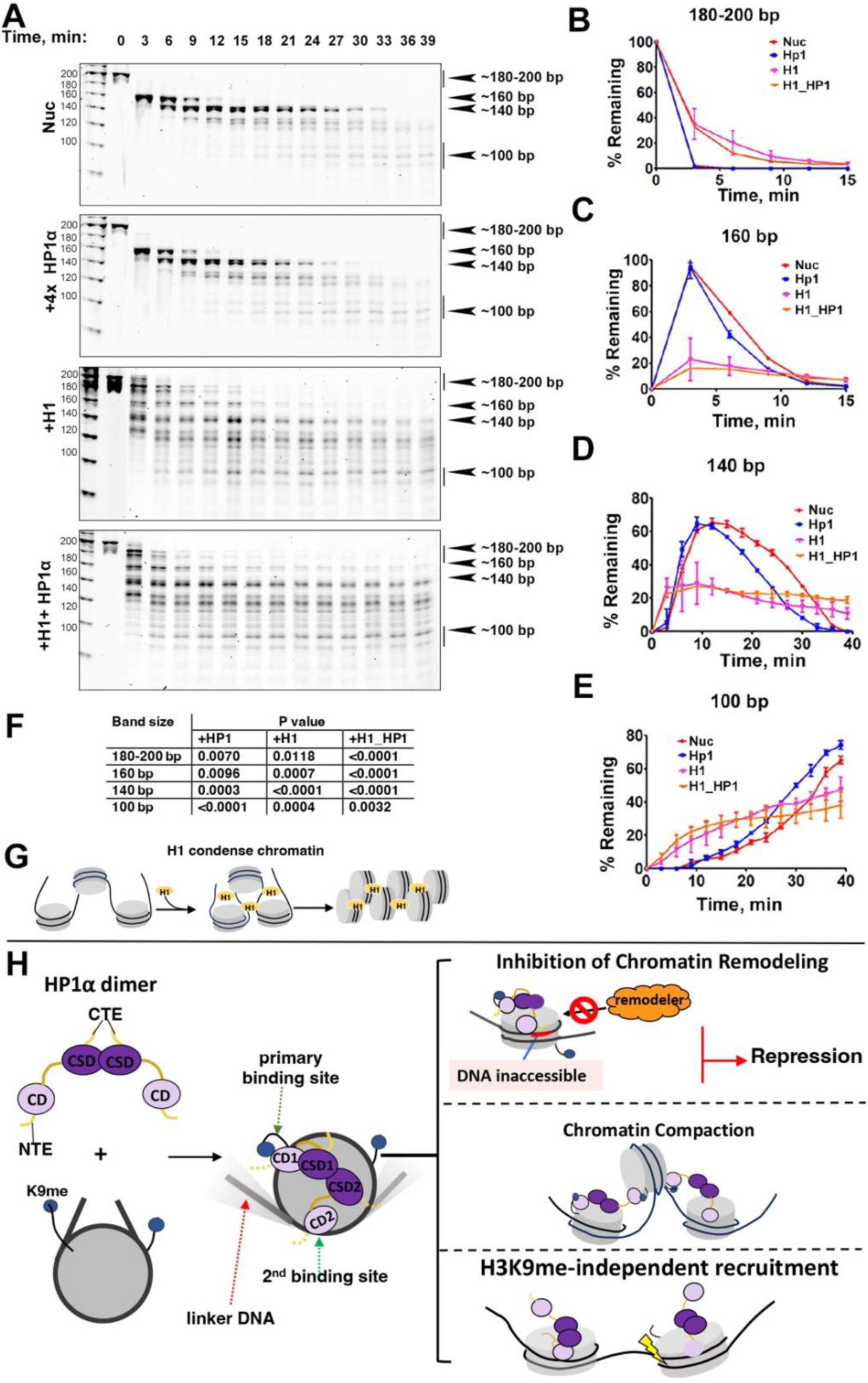
The effects of HP1⍺-binding on nucleosome DNA accessibility. (**A**) Representative acrylamide gels of the MNase assay with no HP1*a*, 4x excess of HP1⍺, 1x H1, and 1x H1+ 4x HP1⍺, respectively. The digestion products of different sizes (180-200, 160, 140 and 100-bp) are labeled. (B) Quantification of the 180-200 bp DNA products shown in (A), representing the fraction of digested nucleosome DNA as a function of time. Data are mean ± SD, n = 3. (C) Quantification of the 160 bp DNA products shown in (A). Data are mean ± SD, n = 3. (D) Quantification of the 140-145 bp DNA product shown in (A). Data are mean ± SD, n = 3. (E) Quantification of the 100 bp DNA products shown in (A). Data are mean ± SD, n = 3. (F) P values of the experiments shown in (B), (C), (D), and (E). (G) Schematic of the model for H1-mediated chromatin compaction. (H) Schematic of the proposed model of HP1⍺-mediated gene repression, heterochromatin maintenance, and H3K9me-independent role in DNA damaged response.

We then explored how HP1⍺ influences MNase digestion of K9me3-H2A.Z nucleosomes. When a 4-fold excess of HP1⍺ (equivalent to two HP1⍺ dimers) was introduced, we observed accelerated digestion over time, particularly for two DNA bands ranging in size from 140 to 147-bp (see **Fig 6D**). This observation aligns with our cryo-EM model’s prediction that HP1⍺ binding enhances the mobility and accessibility of terminal DNAs. Intriguingly, at later time points, we noticed a reverse trend. In the HP1⍺-H2A.Z nucleosome reaction, DNA bands approximately 100 bp in size persisted while they continued to degrade in the control reaction (see **Fig 6E**). A 100 bp product corresponds to the digestion of the 601 Widom sequence at or near SHL-4.5, which is the CD2 binding site revealed by our cryo-EM model. These MNase results collectively suggest that HP1*a* enhances terminal DNA unwrapping while also stabilizing and protecting internal DNA.

## Discussion

### Implications for HP1⍺-mediated transcription repression

In this study, we employed a combination of cryo-EM, MDFF, and XL-MS to obtain a structural model of human HP1⍺ dimer bound to an H2A.Z nucleosome. Our model illustrates how the HP1⍺ dimer asymmetrically engages nucleosomes through extensive interactions between the two CDs and the nucleosome surface. An intriguing discovery from our study is the identification of a second binding site for HP1⍺ on histone H2B, which is independent of H3K9 methylations. Our model also reveals that the function of the HP1⍺ CSD is primarily to mediate HP1 dimerization, without directly engaging in interactions with the nucleosome. Our biochemical data demonstrates that these two CD bindings have contrasting effects on nucleosome DNA accessibility. These findings carry significant implications for our understanding of the multifaceted role of HP1⍺ in shaping heterochromatin and mediating gene expression.

We propose that HP1 binding modulates the structure-dynamics of the nucleosome, resulting in enhanced terminal DNA flexibility while simultaneously protecting internal DNA. Given that many ATP-dependent chromatin remodelers rely on the structural features such as the acidic patch to interact with nucleosomes ^45^, it is conceivable that HP1 binding inevitably compete with chromatin remodelers’ actions on chromatin. Additionally, the binding of HP1 CD domain to histone H2B near SHL4 may further impeded remodelers on nucleosome and their processive DNA translocation activity (**Fig 6H**).

### Implications for HP1⍺-mediated chromatin compaction and maintenance

Linker histone H1, a vital chromatin protein in higher organisms, is renowned for its role in promoting the assembly of higher-order chromatin structures and stabilizing chromatin fibers. Functioning by binding to linker DNAs, H1 forms a stem-like structure with reduced entry-exit angles termed a chromatosome, thereby contributing to the formation of chromatin fibers ^46^ (**Fig 6G)**. Our data suggests that HP1⍺ employs a unique mechanism to facilitate chromatin folding. Instead of constraining entry-exit DNA angles, HP1⍺ increases the flexibility of terminal DNA on nucleosomes, an unexpected outcome resulting from interactions at the primary binding site on histones H3 and H2A. Consequently, in contrast to the regular helical structure observed in *in-vitro* reconstituted H1-chromatin fibers ^47^, we propose that in HP1⍺-containing chromatin, fibers adopt compact yet irregular structures. Our proposition suggests that neighboring nucleosomes are brought into proximity by HP1⍺ dimers through various interactions involving the chromodomains and the two nucleosome binding sites. This model aligns with the idea that the modular nature of HP1 allows the formation of different types of HP1-nucleosome complexes ^48^.

We further propose that the flexible linker DNA enables a higher degree of twisting of linker DNAs between two adjacent nucleosomes, facilitating the formation of compact yet irregular heterochromatin. Although flexible linker DNA may lead to potential higher entropy, HP1-mediated chromatin compaction can also restrict overall nucleosome mobility in the heterochromatin region, compensating for the increase in entropy. Importantly, our model suggests that HP1-heterochromatin is likely polymorphic in nature, aligning closely with recent studies demonstrating that human mitotic chromosomes consist predominantly of irregularly folded chromatin fibers. ^49–51^.

### Chromatin-binding independent of H3K9 methylation

HP1 family proteins are versatile epigenetic regulators with functions outside of heterochromatin maintenance and transcription repression. Early studies indicated a role of HP1 in DNA damage response (DDR) ^52,53^. While many aspects of this enigmatic but crucial role of HP1 in DDR remain unknown, *in-vivo* studies showed that all three HP1 isoforms are efficiently recruited to DNA damage sites in human cells ^54^. This response to DNA damage requires the CSD of HP1, but is independent of H3K9 trimethylation ^54^. The structure presented in our study unveils a novel nucleosome-binding site on the nucleosome surface near histone H2B, away from both copies of the methylated H3 tails. Therefore, our structure provides a rational explanation for how HP1 can be recruited to DNA damage sites in an H3K9me-independent manner.

### The implication for the role of HP1 CSD

CSD-mediated homodimerization has been extensively studied and well characterized. The CSD-CSD dimer, along with the CTE, creates a hydrophobic binding site and is believed to facilitate HP1’s interaction with many non-histone chromosomal proteins containing PXVXL or related motifs (where X denotes any amino acids) ^55,56^. In our model, the CSD-CSD hydrophobic binding site is oriented towards the nucleosome surface but does not directly interact with any histones. Additionally, given that H3 is the sole histone containing a PXVXL sequence at its *a* N helix, it is unlikely that the CSD-CSD hydrophobic pocket makes direct contact with H3 *a* N helix. Our model also suggests that when both CDs are engaged with the same nucleosome, the HP1⍺ dimer may not be capable of binding to PXVXL-containing proteins through CSD-CSD region. However, since our model supports the idea that the modular structure of HP1 allows it to establish various interactions within itself and to form different types of HP1-nucleosome interactions ^48^, it is reasonable to assume that the CSD-CSD hydrophobic binding site will remain accessible for binding partners where only one CD in the HP1 dimer contacts with nucleosome. Future studies will be necessary to test these hypotheses and to gain further insights into these interactions.

## Acknowledgements

We thank the staff in the Laboratory for BioMolecular Structure (LBMS) for help in data collection. LBMS is supported by the DOE Office of Biological and Environmental Research (KP1607011). This work was partially supported by:

NIH grant 1R35GM133611 to D.T.

NSF grant 1942049 to D.T.

Access to the instrumentation was supported by the following grants:

NIH grant U24 GM116790 to M.Y. and D.T.

NIH grant 1S10OD012272-01A1 to H.L.

## Author contributions

V.Sokolova., T.L. and G.L. prepared the samples for EM study; K.D. collected the cryo-EM data; V.Sokolova. performed the biochemical experiment and analysis; D.T. performed cryo-EM image analysis; M.B. performed the AUC experiments and data analysis; J.K. and J.V. performed model building under the supervision of A.S.; V. Svetlov performed the XL-MS experiments under the supervision of E.N.; D.T. oversaw the project and wrote the manuscript with help from all authors.

## Data and materials availability

The data that support the findings of this study are openly available in the Electron Microscopy Database (https://www.ebi.ac.uk/pdbe/emdb) and the Protein Data Bank (https://www.rcsb.org/). EM maps are deposited at the Electron Microscopy Database under accession code EMD-42774 for the HP1⍺-H2A.Z nucleosome complex (map1), EMD-42690 for focused refined HP1⍺ in complex with partial H2A.Z nucleosome (map2), EMD-42692 for focused refined K9me3_H2A.Z nucleosome (map3), and EMD-42773 for the free K9me3_H2A.Z nucleosome. The protein coordinate of the HP1⍺-H2A.Z nucleosome complex presented in the paper is deposited at PDB ID 8UXQ).

## Material and Methods

### Expression and purification of HP1*a*

GST-tag human HP1⍺ was a kind gift from Naoko Tanese (Addgene plasmid # 24074; http://n2t.net/addgene:24074; RRID:Addgene_24074). GST-tag HP1⍺ΔC construct was generated by site-direct mutagenesis kit (NEB) by deleting the amino acids (number 169 –181) of the protein. GST-HP1 and GST-HP1ΔC were expressed in BL21 (DE3) E. coli cells. Both proteins were purified using the same protocol as follows. Cells were lysed by sonication in buffer containing 50 mM Tris-HCl pH7.5, 400 mM NaCl, 10% glycerol supplemented with protease inhibitors and 1mM Dithiothreitol (DTT). Cell lysate was subjected to batched affinity chromatography purification using glutathione sepharose (Cytiva). HP1 proteins was released from the GST-tag by overnight thrombin digestion followed by purification on a 50HQ 10×100 column (Applied Biosystems) in a continuous gradient of 50 –1000 mM NaCl. Peak fractions were pooled and further purified by gel filtration with a Superdex 200 increase 10/300 GL column (GE Healthcare). Peak fractions were pooled and concentrated to 11 mg/ml.

### Nucleosome reconstitutions

The plasmid with twelve tandem repeats of 208-bp 601 Widom sequence was a kind gift from Dr. Ed Luk. Large-scale plasmids were purified as previously described ^57^. Restriction enzyme ScaI was used to excise the plasmids to generate single repeat of 208-bp segment. DNA fragments were further purified by polyethylene glycol precipitation and MonoQ anion exchange chromatography.

The sequence for the 208-bp Widom 601 DNA repeat is as follows: ACTTATGTGATGGACCCTATACGCGGCCGCCCTGGAGAATCCCGGTGCCGAGGCCGCTCAATTGGTCGTAGACAGCTCTAGCACCGCTTAAACGCACGTACGCGCTGTCCCCCGCGTTTTAACCGCCAAGGGGATTACTCCCTAGTCTCCAGGCACGTGTCAGATATATACATCCTGTGCATGTATTGAACAGCGACCTTGCCGGAGT

Canonical *Xenopus* laevis histones H2B and H4 were expressed in BL21 (DE3) pLysS *E. coli* cells and purified as previously described ^57^. Mouse H2A.Z.1 gene in pIND-EGFP was a kind gift from from Danny Rangasamy (Addgene plasmid # 15770; http://n2t.net/addgene:15770; RRID:Addgene_15770). It was re-cloned into pET-LIC expression vector. The protein was expressed in BL21 (DE3) *E. coli* cells and purified using the same procedure as the canonical histones. To produce histone H3 containing H3K9me3 mimic, a single point mutation K9C was first introduced on *Xenopus laevis* H3. A procedure to install a tri-methyl-Lysine analog (tri-MLA) on C9 was then performed as described ^58^. For some nucleosome preparations, the same modified H3 was purchased from The Histone Source at Colorado State University.

Histone octamers containing the abovementioned histones were produced *in vitro* using salt dialysis as previously described ^57^. Briefly, equal molar of each histone was mixed and incubated for 2 h in unfolding buffer (7 M guanidine HCl, 20 mM Tris, pH 7.5, and 10 mM DTT) followed by dialysis against at least three changes of refolding buffer (10 mM Tris, pH 7.5, 1 mM EDTA, pH 8.0, 2 M NaCl and 1 mM DTT) at 4°C. Octamer was concentrated and purified by gel filtration using a Superdex200 increase 10/300 GL column. Mono-nucleosomes were reconstituted by mixing the octamer with 208bp 601 Widom sequence DNA in equal molar ratio in high-salt buffer (10 mM Tris, pH 8.0, 2 mM EDTA, 2 M NaCl and 2 mM 2-Mercaptoethanol (βME) followed by overnight dialysis into low salt buffer (10 mM Tris, pH 8.0, 2 mM EDTA, 5 mM NaCl and 2 mM βME) as described ^57^.

### SV-AUC

SV-AUC studies were conducted in a Beckman XL-I using the absorption optics to scan cells assembled with a double sector charcoal EPON centerpiece and sapphire glass windows inserted into a AN-60 Ti rotor. All samples were exchanged into buffer containing 10 mM HEPES, 75 mM NaCl, and 0.5 mM TCEP at pH 7.5. The SV-AUC runs were conducted at 4°C and 48,000 rpm for HP1⍺ and 38,000 for the HP1⍺-nucleosome complexes. For HP1⍺, samples with protein concentrations of 0.52, 1.82, and 4.15 μM were scanned at 230 nm. HP1⍺ samples with protein concentrations of 3.38, 11.16 and 23.57 μM were scanned at 280 nm. The nucleosome and HP1⍺-nucleosome complex samples were scanned at the absorption maximum of DNA, 260 nm. Nucleosome at a constant concentration of 0.16 μM was titrated with increasing concentrations of HP1⍺ ranging from 0.1 to 23.73 μM. The data point at 0.01μM is nucleosome without added HP1⍺.

A value of vˉ = 0.7282 was calculated from the sequence of HP1⍺ using the program SEDNTERP ^59^. A measured value of vˉ = 0.65 for a nucleosome was used to analyze the HP1⍺-nucleosome complexes ^60^. The sedimentation parameters were corrected to S(20,w) using values of ρ = 1.00393 and η = 1.5926 at 4°C calculated with Sedenterp from the buffer composition. Continuous distribution component analysis, c(s), was conducted using the program Sedfit ^61,62^ to deconvolute the species present in a solution. The software Prism version 10.0.1 was used to plot the c(s) distributions and the dependence of S(20,w) on HP1⍺ and HP1⍺-nucleosome complex concentration. The HP1⍺-nucleosome titration was fit to the Hill equation with a function that explicitly calculates the free concentration of the HP1⍺ ligand.

### MNase assays

For the digestion assay with MNase, 340 nM nucleosome was subjected to digestion with 0.75U of MNase (Roche) in a 65 µl reaction in MNase buffer (10 mM Tris pH7.4, 50 mM NaCl, 2 mM CaCl_2_) in the presence of 4-fold excess of HP1⍺ at 37°C. For experiments containing linker histone H1.0 (NEB # M2501S), 1:1 ratio of H1:nucleosome was used. Nucleosomes with or without HP1⍺ and/or H1.0 were mixed and incubated for 30min prior to MNase digestion. Samples (4.5 µl) were collected every 3 min. 8 µl stop/deproteinization buffer (10 mM Tris–HCl, pH 8.0, 0.6% SDS, 40 mM EDTA, 0.1 mg/ml proteinase K) was added to quench the reaction followed by 1 hour incubation at 55°C. The samples were resolved on 8% (19:1 Acrylamide/Bis) Native-PAGE (at 100V, 120 min). The gel was stained with SYBR-Gold and imaged with Typhoon imager (Cytiva). Images were analyzed using ImageJ software. Intensities for fragments with sizes of 180-200, 160, 140, and 100 bp were calculated. Two-way Anova test was used to determine the level of significant difference (p≤0.05). Two-way Anova test and graphical representation was done using Prism 5 software.

### Nucleosome Binding assay

HP1⍺ and HP1⍺ΔC was dialyzed for 3 hours in binding buffer (10 mM Tris, pH 8. 25 mM NaCl, 2 mM EDTA, 1 mM DDT) separately prior to binding reaction. 250nM in 15 µl were mixed with increasing amount of HP1⍺. The mixture was incubated on ice for 30 min. The reactions were there resolved on 4% Native-PAGE (100 V, 90 min) and stained with SYBR-Gold before imaging by FluorChem8900 Imager (Alpha Innotech).

### Assembly of the HP1*a* –H2A.Z nucleosome complex for cryo-EM study

Three Gradient Fixation (GraFix) experiments with three different protein preparations were conducted to produce the samples used for the cryo-EM study. In two Grafix experiments 1.2 µM of nucleosome was mixed with 75-fold excess of HP1⍺ (90 µM) in 500 µl reaction in binding buffer A (10 mM Tris, pH 8. 25 mM NaCl, 2 mM EDTA, 1mM DDT). In the third experiment, similar condition was applied except that 32-fold excess of HP1*a* (38.4 µM) and buffer containing 75 mM NaCl were used. In each experiment, HP1*a* –nucleosomes mixture was incubated on ice for 30 min and then concentrated to 200ul before subjected to GraFix ^63^ for separation and cross-linking. The continuous density gradient was formed by mixing two buffer solutions with a Gradient Master (BioComp). The top buffer contains 10 mM HEPES, 20 mM NaCl, 1mM DTT and 5% sucrose, while the bottom buffer contains 10 mM HEPES, 20 mM NaCl, 1 mM DTT, 25% sucrose and 0.2% glutaraldehyde. Ultracentrifugation was carried out at 4℃ in a SW41 rotor (Beckmann) for 16 hours with speeds of 27,000 rpm. Following centrifugation, the gradients are fractionated. The optical density of each fraction was measured with a Nanodrop Spectrophotometer (Thermo Fisher). Peak fractions were further examined by Native-PAGE gels and by negative-stained EM to identify the complex. Complex formation was observed in all three experiments, within a similar pattern of complex separation from free nucleosomes. Fractions containing the complex were pooled, dialyzed into storage buffer (10 mM HEPES pH 8, 20 mM NaCl, 1 mM DTT) and concentrated to 2-3.8 µM.

### Cryo-EM sample vitrification

Cryo-EM grids were prepared using Vitrobot Mark IV (FEI Company) under 8℃ and 100% humidity. Aliquots of 3.5 µl of the HP1⍺-H2A.Z nucleosome complex or the free K9me3-H2A.Z nucleosome were applied to glow-discharged QUANTIFOIL grids (R1.2/1.3 – 400 mesh), blotted for 4 to 5 seconds and plunged into liquid ethane cooled by liquid nitrogen. Grids were stored in liquid nitrogen until they were imaged.

### Cryo-EM data collection

Grid screening was done using the Talos Arctica microscope (FEI) at the cryo-EM facility in Stony Brook University to identify suitable grids for data collection. For the HP1⍺-nucleosome samples, multiple datasets were collected at the University of Virginia Molecular Electron Microscopy Core using the Titan Krios electron microscope (FEI Company) operating at 300 kV with a nominal magnification of 81,000X, giving a pixel size of 1.08 Å at the specimen level. Movies were recorded using a K3 direct detector (Gatan company) in counting mode using EPU software, with the Bioquantum energy filter operating at zero loss frequency 10 eV. Defocus values range from – 1.0 to –2.25 µm. Each movie was dose-fractionated to 40 frames with a dose rate of ∼1.25 e/Å^2^/sec. Total dose per micrograph is 50 e/ Å^2^ (**Extended Data Table 1**).

For the K9me3-H2A.Z nucleosome sample, one dataset was collected at the Laboratory of Biomolecular Structure at Brookhaven National Laboratory (BNL) using the Titan Krios electron microscope (FEI Company) operating at 300 kV with a nominal magnification of 105,000X giving a pixel size of 0.825 Å at the specimen level. Movies were recorded using a K3 direct detector (Gatan company) in counting mode using EPU software, with the Bioquantum energy filter operating at zero loss frequency 15 eV. Defocus values range from –1.0 to –2.5 µm. Each movie was dose-fractionated to 40 frames with total dose per micrograph being 40 e/ Å^2^ (**Extended Data Table 1**).

### Image analysis

For the HP1⍺-nucleosome datasets, frames were aligned and summed using the motion correction software implemented within RELION ^64^. The CTF parameters were estimated using CTFFIND4 ^65^. Particle-picking was carried out using Topaz neural-network picking ^66^. Both particle-picking and 2D classification were done in cryoSPARC ^67^, while the rest of image processing steps were carried out in RELION. Specifically, bad particles were removed through multiple rounds of 2D classifications, resulting in a data set of 1.3-millions of particles, which were then subjected to 3D classification in RELION. Good 2D classes representing different views of the complex were selected and used for the Ab-Initio reconstruction in cryoSPARC. The subsequent low-resolution map was used as the initial model for 3D classification in RELION. After three rounds of 3D classifications, the best class containing 74,257 particles was selected and subjected to consensus 3D refinement. The particles were then subjected to Bayesian Polishing and postprocessing, yielding map with an average resolution of 4.1 Å. The global resolution of the map was estimated using the gold standard Fourier Shell Correlation (FSC) 0.143 criterion with automatic B factor determined in RELION. Local resolution was estimated in RELION.

To improve the resolution of the HP1 region, we performed signal subtraction and focused refinement in RELION. Briefly, the consensus map was used to generate two overlapping masks using the Volume eraser tool in UCSF ChimeraX. One mask contains only the nucleosome region, while the second mask contains HP1 density and most of the nucleosome (mask 2 and mask 3 in **Extended Data** Fig 3C). Multibody refinement was then performed using these two masks. The subsequent volumes representing the two body were used as new masks to perform signal subtraction and focus-refinement, which resulted in improved resolution and map feature of the two bodies.

Image processing of the K9me3-H2A.Z nucleosome dataset was performed using cryoSPARC ^67^. Briefly, a dataset containing 454,841 particles were generated after Topaz neural-network particle picking followed by two-rounds of 2D classification. This dataset was then subjected to additional rounds of 3D heterogeneous refinement, which resulted in a final subset of 186,172 particles. Non-uniform refinement was performed with this subset to generate the final map (**Extended Data** Fig 5).

All conversions between RELION and CryoSPARC were performed using D. Asarnow’s pyem script (personal communication; https://github.com/asarnow/pyem). The generation of figures featuring images related to the structural model was carried out using UCSF ChimeraX ^68^ and Coot ^69^. Movies were created using UCSF ChimeraX.

### Model building and refinement

The specific refinement protocol applied to the HP1 model involves iteratively applying Cascade Molecular Dynamics Flexible Fitting (cMDFF) ^28^, Modelling Employing Limited Data (MELD) ^29^, and Molecular Dynamics Flexible Fitting (MDFF) ^28^ to optimize the atomic coordinates of the model to match the experimental density map obtained through Cryo-EM (**Extended Data** Fig 7). This process is described in more detail below. The cross-correlation coefficient, a commonly used metric in the field of structural biology, is employed as a measure of the similarity between the experimental density map and the atomic model. This metric is used to evaluate the quality of the refined model.

To generate an initial model for refinement, a modified H2A.Z nucleosome coordinate (combining the histone octamer of PDB 1F66 with the Widom 601 DNA from PDB 6FQ5) and the Alphafold model of full-length human HP1*a* (AF-P45973) was used. The H2A.Z nucleosome placement was determined by rigid-body fitting in UCSF ChimeraX. For HP1*a*, we extracted the different domains as individual molecules and used UCSF ChimeraX to perform rigid-body docking to determine their initial placements. The initial positions of CDs and CSD-CSD were determined by docking these molecules separately in the focused refined map of the HP1⍺-nucleosome complex (map 2 in **Extended Data** Fig 3 **&4**), guided by the visible secondary structures. Using this strategy, the positions of two CD2 and the CSD-CSD dimer can be unambiguously determined. The flexible regions of NTE, HR, and CSD were excluded during this process. The relative conformations between the two CSD domains, similar to the CSD-CSD dimer conformation in the crystal structures of Drosophila CSD-CSD dimers ^33^, were preserved during the manual rigid-body fitting. Once the optimal placements of the CDs and CSD-CSD domains were identified, the flexible loops (NTE, HR and CTE) were added and reconnected to form a full-length HP1⍺. The reconnection was done using VMD’s psfgen tool which takes in the necessary topology files and generates the appropriate structure files. Specifically, patches were applied between the CD / CSD domains and the respective NTE, HR, and CTE domains followed by minimization to alleviate nonphysical bonds and structural features. This process yielded an initial structure of the HP1 dimer-H2A.Z nucleosome complex (**Extended Data** Fig 5B)

#### Equilibrium MD

The initial model was then subjected to a minimization and 50 ns equilibration simulation using GPU-accelerated NAMD3 software, implementing CHARMM36 force fields and a TIP3 water model parameterization for the protein / nucleic and solvent components.

#### MDFF/cMDFF

The subsequent equilibrated structure was fitted to a series of four density maps each with increased resolution using the cMDFF protocol (**Extended Data** Fig. 5B). The maps were created by smoothing the original density map, obtained from Cryo-EM data, by applying a Gaussian blur with a half-width parameter σ. The first map applied a Gaussian blur with σ = 6 Å; each subsequent map was smoothed by 2 Å less than the previous map until the final map which corresponded to the original Cryo-EM map resolution. Each MDFF fitting simulation was run for 50 ns using NAMD3 software and CHARMM36 force fields and was restricted to the backbone atoms. The fit of the model was evaluated using VMD timeline analysis, where a cross-correlation coefficient (CCC) was calculated at each time step along the trajectory to measure the degree of fitness of the model with respect to the density of interest.

#### MELD

The MELD protocol was integrated into the computational workflow in conjunction with cMDFF and MDFF, as illustrated in (**Extended Data** Fig. 7 **A&B).** This design is an extension of the CryoFold algorithm, a multiphysics approach that generates equilibrium ensembles of biomolecular structures from cryo-EM data ^70^, leading to a comprehensive sampling of the conformational landscape of the complex. The incorporation of MELD allowed for the implementation of precise restraint applications, which are not feasible using standard restrained MD methods. These restraints were applied as flat-bottom harmonic potentials, and their satisfaction resulted in no additional contribution to the energy function, i.e. they fell within the flat-bottom region. However, for restrictions that fell outside this region, energy penalties were applied, which acted as a redirecting force guiding the system towards low-energy basins.

The MELD simulations used a Replica Exchange Molecular Dynamics (REMD) sampling method with a 16-replica parameterization, a temperature range of 300-380K, a GBNeck2 implicit solvent model, and a simulation time of 250 ns. A linear temperature ramp was applied throughout the one-dimensional Hamiltonian exchange ladder starting from replica 1 and finishing at replica 6. Additionally, distance restraints were applied linearly throughout the ladder starting at replica 7 and finishing at replica 16 with a force constant of 250 kJ / (mol * nm2). Specifically, cartesian restraints were enforced in regions with a high map correlation determined from earlier cMDFF results. These regions were assigned confidence values of 100% in each and every replica (1-16). Distance restraints between alpha carbons were applied throughout the remaining domains starting at replica 7 and finishing at replica 16. These restraints were applied with varying confidence values ranging from 50-90% based on external experimental information. This approach resulted in an ensemble of the lowest free energy structures consistent with experimental knowledge. The bsc1 and ff14SB forcefields in AMBER were used to generate the appropriate topology and parameter file for MELD simulations.

These procedures resulted in a marked improvement in the CCC, however, certain regions, particularly flexible loop regions throughout the model remained positionally ambiguous, necessitating further refinement and sampling of the conformational space. To preserve the accuracy of the mapping obtained from cMDFF, high-energy Cartesian restraints were applied to the alpha carbons in regions that displayed high map correlation from cMDFF. These restraints penalized any deviation of more than 0.2 nm from the initial Cartesian coordinates, thus restricting their deviation from the initial positions. Additionally, the restraints were lifted from flexible loop regions to optimize the exploration of the conformational space, reflecting the likelihood function utilized throughout the MELD simulations. The constrains applied to different regions of the complex were indicated in **Extended Data** Fig 7C. MELD simulations reveal various conformations of the flexible loop regions, notably the H3 tail (**Extended Data** Fig. 7B), that better fit the density.

### XL-MS and data analysis

Bissulfosuccinimidyl suberate (BS3) Crosslink test was performed to test and analyze stable amide bonds. K9me3-H2A.Z nucleosomes (2.44 µM) and 4-fold access of purified recombinant human HP1α (9.76 µM) were mixed in a final volume of 50 µl reaction and dialyzed into binding buffer [75 mM NaCl, 10 mM HEPES, 0.4 mM 2-Mercaptoethanol and 1 mM DTT] at 4°C for 3 hours. Then, fresh 2 mM DTT was added. BS3 crosslinker (250 µM and 1 mM respectively) was added to the HP1α control and the HP1α-nucleosome complex and the reactions were incubated at 25°C for 45 minutes. After the incubation, samples were quenched with 50 mM Tris-HCl pH7.5 for 15 minutes at room temperature. The samples were flash frozen in liquid nitrogen then stored in – 80°C freezer until XL-MS analysis.

Frozen samples were thawed and were separated using SDS-PAGE (200 V, Thermo Fisher NP0050, NP0335BOX) and stained with GelCode Blue (Thermo Fisher 24592). Supershifted bands were excised from the gel, destained twice with 100-fold (m/m) excess of 50% acetonitrile (ACN, Millipore Sigma 900667) in 25 mM LC-MS-grade ammonium bicarbonate (ABC, Honeywell Fluka, 40867), 37^0^C, 30 min, with agitation. Destained slices were replaced into 100 µl of 50 mM tris-2-carboxyethyl)phosphine (TCEP, Millipore Sigma, 646547), incubated for 10 min at 60°C, and destained as described above. Destained sliced were incubated in 100% ACN, and air dried for 10 min, before addition of 75 µl of 25 mM ABC, supplemented with 50 ng/µl of MS-grade trypsin (Thermo Fisher 90305) and 0.01% ProteaseMAX Surfactant (Promega, V2071). Digestion was carried out for 120 min at 54^0^C, peptides were collected as described in ProteaseMax application note (Promega, TB373, Rev. 2/15), and purified using Pierce C18 Spin Tips (Thermo Fisher PI84850). Peptide samples were dissolved in 0.1% (v/v) formic acid (FA, Thermo Fisher Scientific, LS118-4) for MS analysis. Peptides were analyzed in an Orbitrap Fusion Lumos mass spectrometer (Thermo Scientific) coupled to an EASY-nLC (Thermo Scientific) liquid chromatography system, with a 2 μm, 500 mm EASY-Spray column (Thermo Scientific, ES903). The peptides were loaded on the column in 100% buffer A (0.1% FA in water), and eluted at 200 nl/min over either linear 140 min gradient 4-40% buffer B (0.1 % FA in ACN, Thermo Scientific 85174), or 195 min concave gradient (4-30%, curve value set at 6). Each full MS scan (R = 60,000) was followed by 20 data-dependent MS2 (R = 15,000) with high-energy collisional dissociation and an isolation window of 2.0 m/z. The normalized collision energy was set to 30%. Notable selection algebra included 4-6 charge precursor isolation window, TopN, [lowest charge then most intense] operators. Monoisotopic precursor selection was enabled and the dynamic exclusion window was set to 30.0 s. Resulting raw files were searched in enumerative and cross-link discovery modes. Enumerative mode was engaged by applying open search implemented in the pFind3 software ^71^ using the fasta file combining sequences of the recombinant targets and the expression host proteome(*E. coli*, Uniprot UP000000625) as the search space. Fasta sequences of proteins identified in the samples in the enumerative mode were combined to form the search space for crosslink discovery by pLink2 ^72^; protein modifications inferred by pFind3 and comprising >0.5% of the total were included as the variable modifications in pLink2 search parameters. pLink2 results were filtered for FDR (<5%), e-value (<1.0E-3), score (<1.0E-2).

### Quantification and statistical analysis

In Figure 6B, 6C, 6D, and 6E, the average values of three biological replicas were shown with the standard deviation of the mean (SD) in each case. In all cases, reproducible results were obtained.

