## Supplemental Figures for "Structural mechanism of HP1α-dependent transcriptional repression and chromatin compaction"

**Extended Data Table 1. Cryo-EM data collection**

| Dataset | | HP1⍺-H2A.Z nucleosome  (EMD-42774)  (PDB ID 8UXQ) | K9me3_H2A.Z nucleosome  (EMD-42773) |
| --- | --- | --- | --- |
| **Data acquisition and processing** |  | |  |
| Microscope  Voltage (kV) | Krios  300kV | | Krios  300kV |
| Detector  Magnification | K3  81,000X | | K3  105,000X |
| Defocus range (µm)  Total dose (e/ Å^2^) | -0.9 to -2.5 µm  50 | | -1 to -2.5 µm  40 |
| Symmetry | C1 | | C1 |
| Initial particle images (no.) | 1,281,469 | | 454,841 |
| Final particle images (no.) | 74,257 | | 186,172 |
| Map resolution (Å) | 4.1 | | 3.5 |
| FSC threshold | 0.143 | | 0.143 |
| Symmetry | C1 | | C1 |
| Map sharpening B-factor (Å) | -120.654 | | -98.126 |

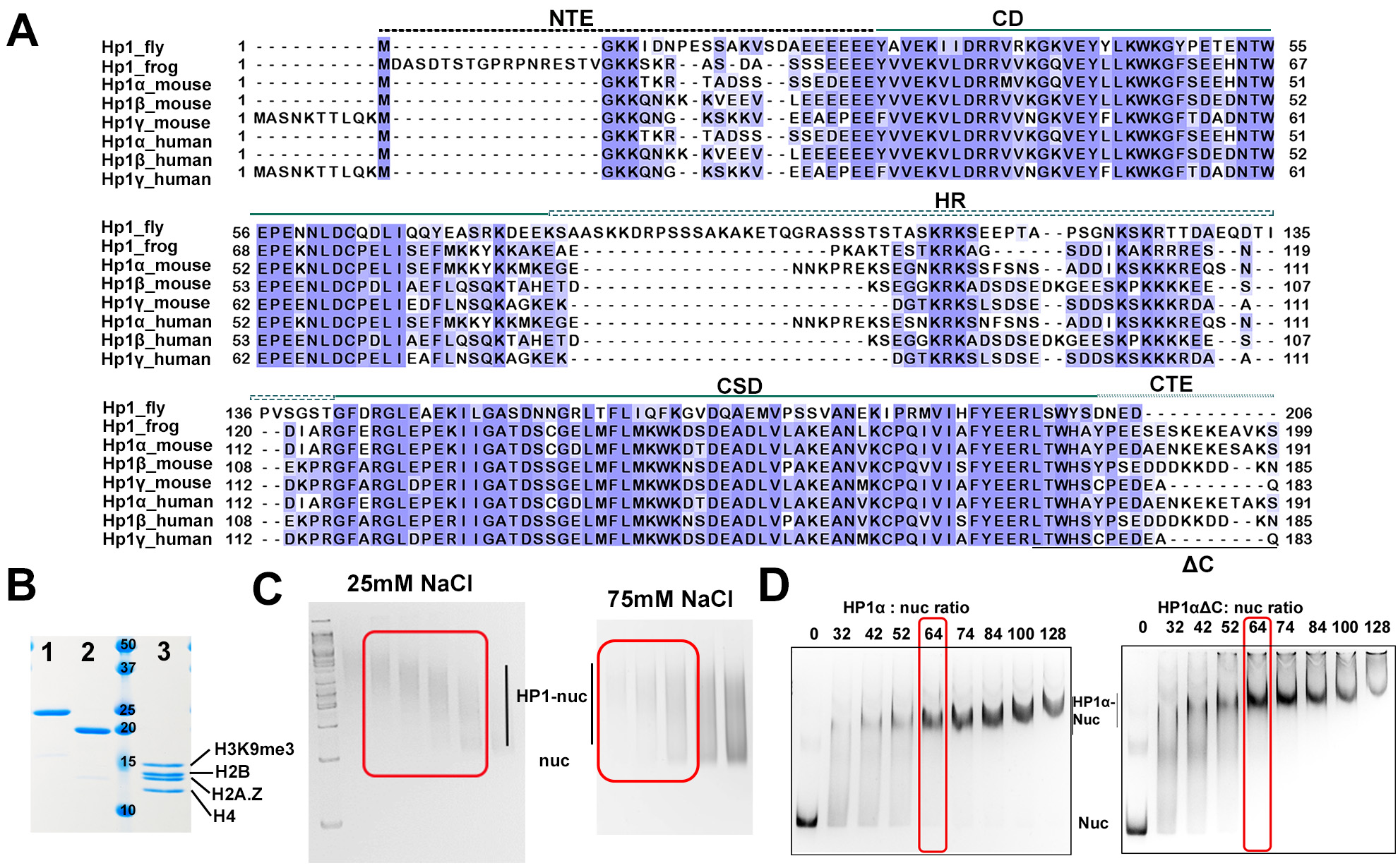

**Extended Data Fig 1. Human HP1⍺ protein preparation and H3K9me3-H2A.Z-nucleosomes binding**

A. Sequence alignment of HP1 family proteins to show the conservation of different domains: N-terminal extension (NTE), Chromo domain (CD), the Hinge Region (HR), the Chromo shadow domain (CSD), and the C-terminal extension (CTE). The region removed in the HP1𝒶𝚫C construct is underlined.
B. SDS-PAGE showing the purified recombinant human HP1𝒶 (lane #1), HP1𝒶𝚫C (lane #2) and all the histones used for reconstituting the H3K9me3-H2A.Z nucleosome (lane #3).
C. Representative native gels of GraFix fractions containing the HP1𝒶-nucleosome complex (highlighted in red box). The left panel represents an experiment conducted at 25 mM NaCl (in HP1-nucleosome binding buffer), and the right panel represents an experiment conducted under 75 mM NaCl.
D. Native gels show that HP1𝒶𝚫C binds to K9me3-H2A.Z nucleosomes (right panel) at the same HP1:nucleosome ratio (red box) used for the HP1𝒶 full-length protein (left panel).

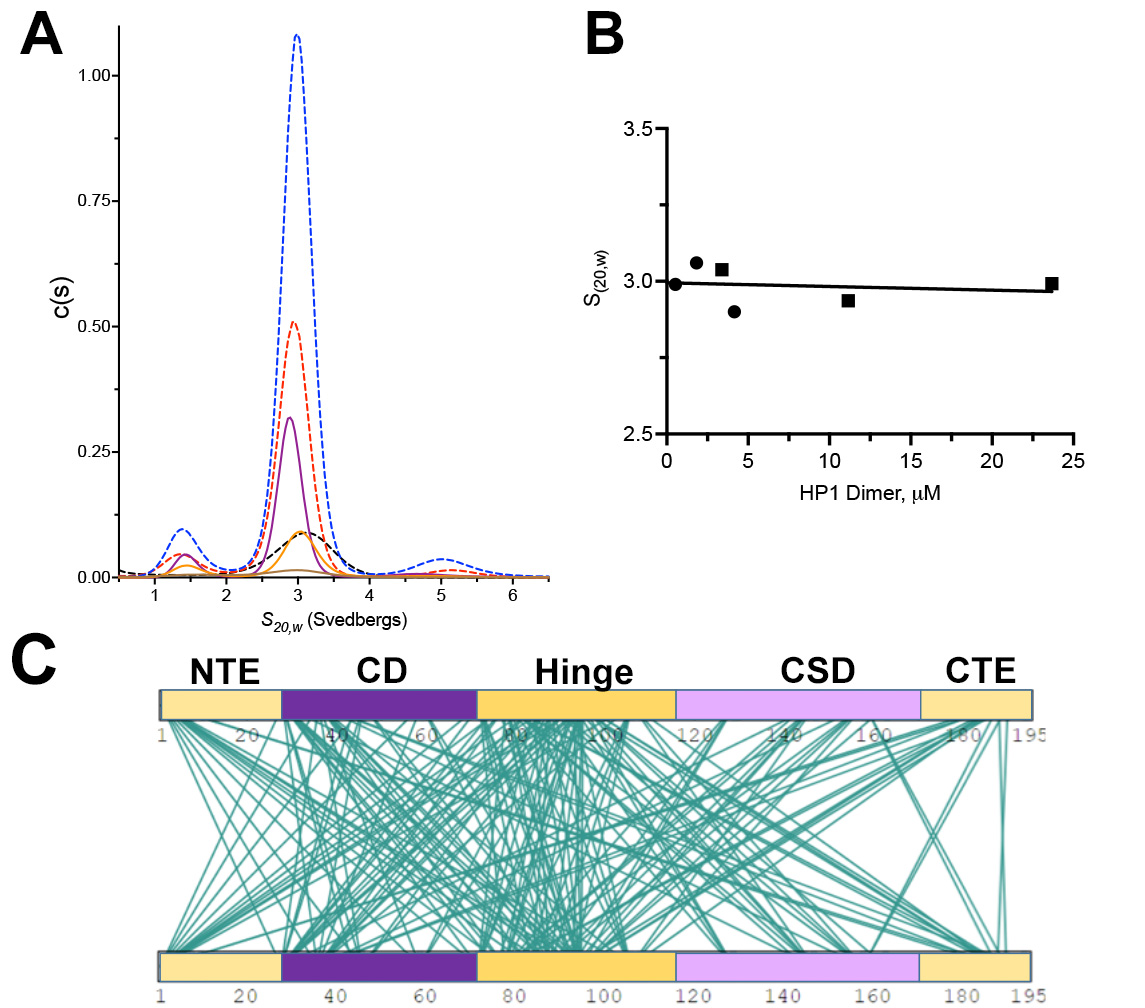

**Extended Data Fig 2. SV-AUC and CL-MS analysis of recombinant HP1𝒶**

A. The c(s) distributions obtained for HP1⍺ as a function of protein concentration. The HP1⍺ at 23.8 (dashed blue), 11.2 (dashed red), and 3.4 (dashed black) μM were scanned at 280 nm. The samples at 0.5 (magenta), 1.8 (orange), and 4.4 (brown) μM were scanned at 230 nm.
B. Summary of the protein concentration dependence of the *S*_20,w_ values determined for the dominant peak (HP1⍺ dimer) of the c(s) distributions shown in (A). The circles depict data obtained by scanning the samples at 280 nm. The squares depict data obtained by scanning the samples at 230 nm. The solid line is the best fit linear regression to the *S*_20,w_ values.
C. Crosslinks within the free HP1𝒶 dimer identified by the XL-MS experiments.

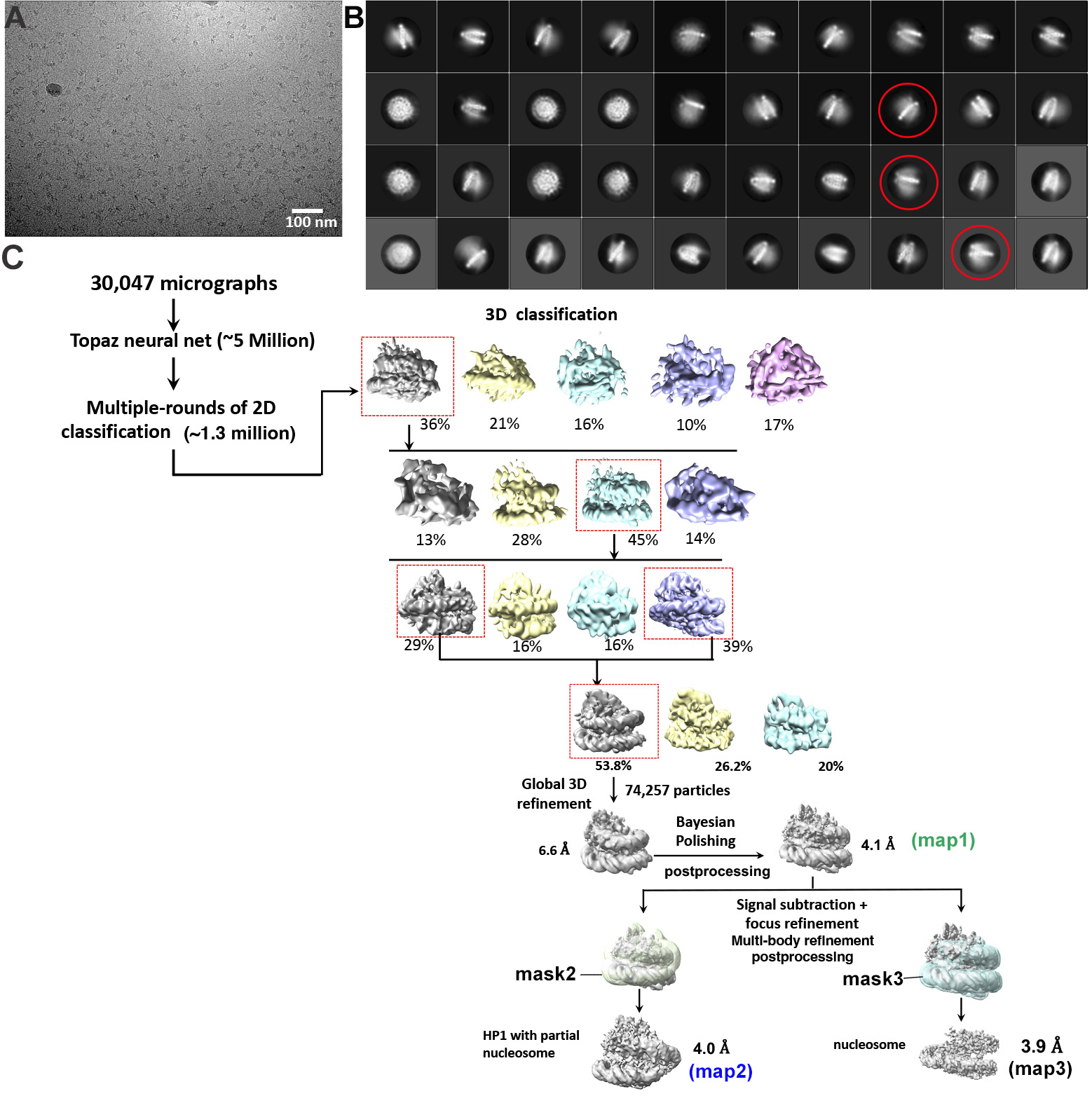

**Extended Data Fig 3. Single-particle cryo-EM data analysis.**

A. A representative motion corrected cryo-EM image of the HP1𝒶- H2A.Z nucleosome complex.
B. Selected 2D classes of the HP1𝒶-H2A.Z nucleosome complex. Classes with densities on both side of the nucleosome are highlighted in red circles.
C. Schematic depicting the single-particle image-processing workflow.

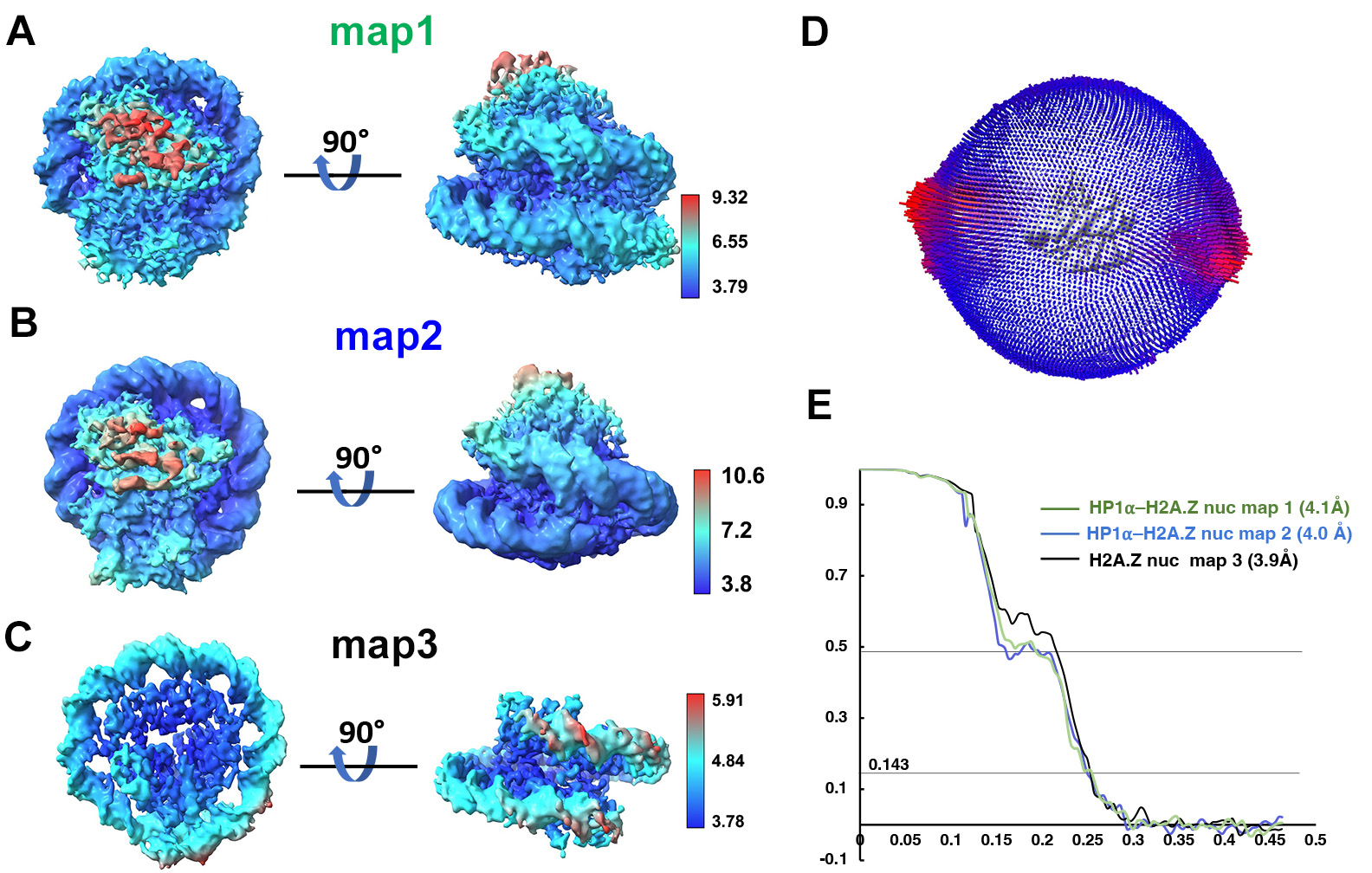
**Extended Data Fig 4. Validation of the cryo-EM maps.**

A. Local resolution of the consensus refined map of HP1𝒶- H2A.Z nucleosome complex (map 1 in Extended Data Fig 3).
B. Local resolution of the signal-subtracted, focused refined map of HP1𝒶- H2A.Z nucleosome complex (map2 in Extended Data Fig 3).
C. Local resolution of the signal-subtracted, focused refined map of the H2A.Z nucleosome in the complex (map3 in Extended Data Fig 3).
D. Angular distribution for the particles in the final dataset used for the consensus refinement.
E. Gold-standard Fourier Shell correlation (FSC) curves to show the average resolutions of the three maps after postprocessing.

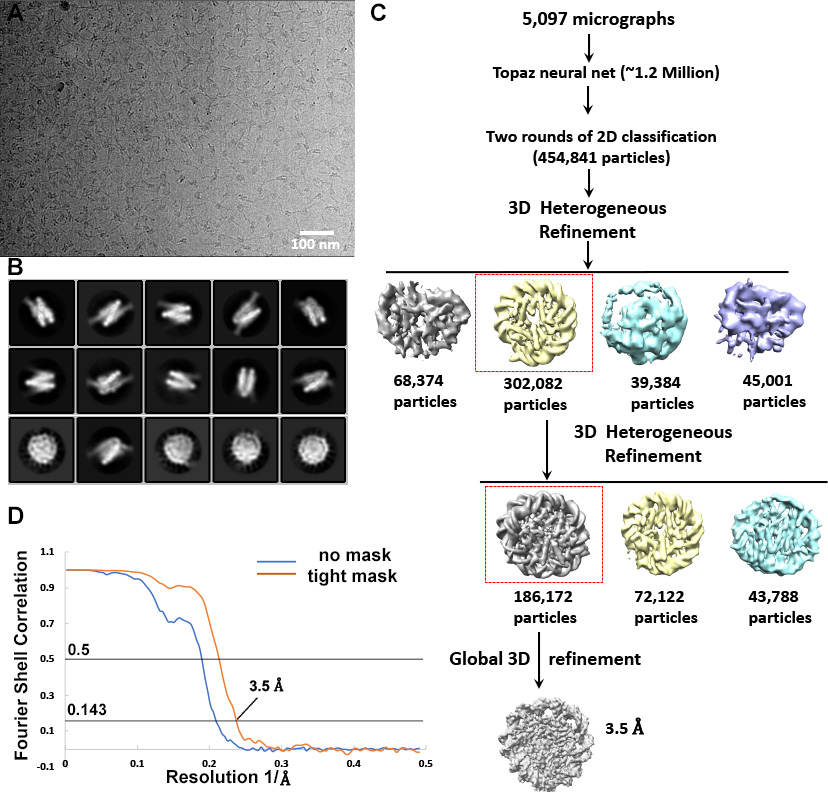

**Extended Data Fig 5. Single-particle cryo-EM data analysis of K9me3- H2A.Z-nucleosome**

A. Selected motion corrected cryo-EM image of K9me3-H2A.Z nucleosomes.
B. Selected 2D classes of K9me3-H2A.Z nucleosomes.
C. Schematic depicting the single-particle image-processing workflow.
D. Gold-standard Fourier Shell correlation (FSC) curve to show the estimated resolution of the consensus refined map of the H3K9me3-H2A.Z nucleosome with and without mask.

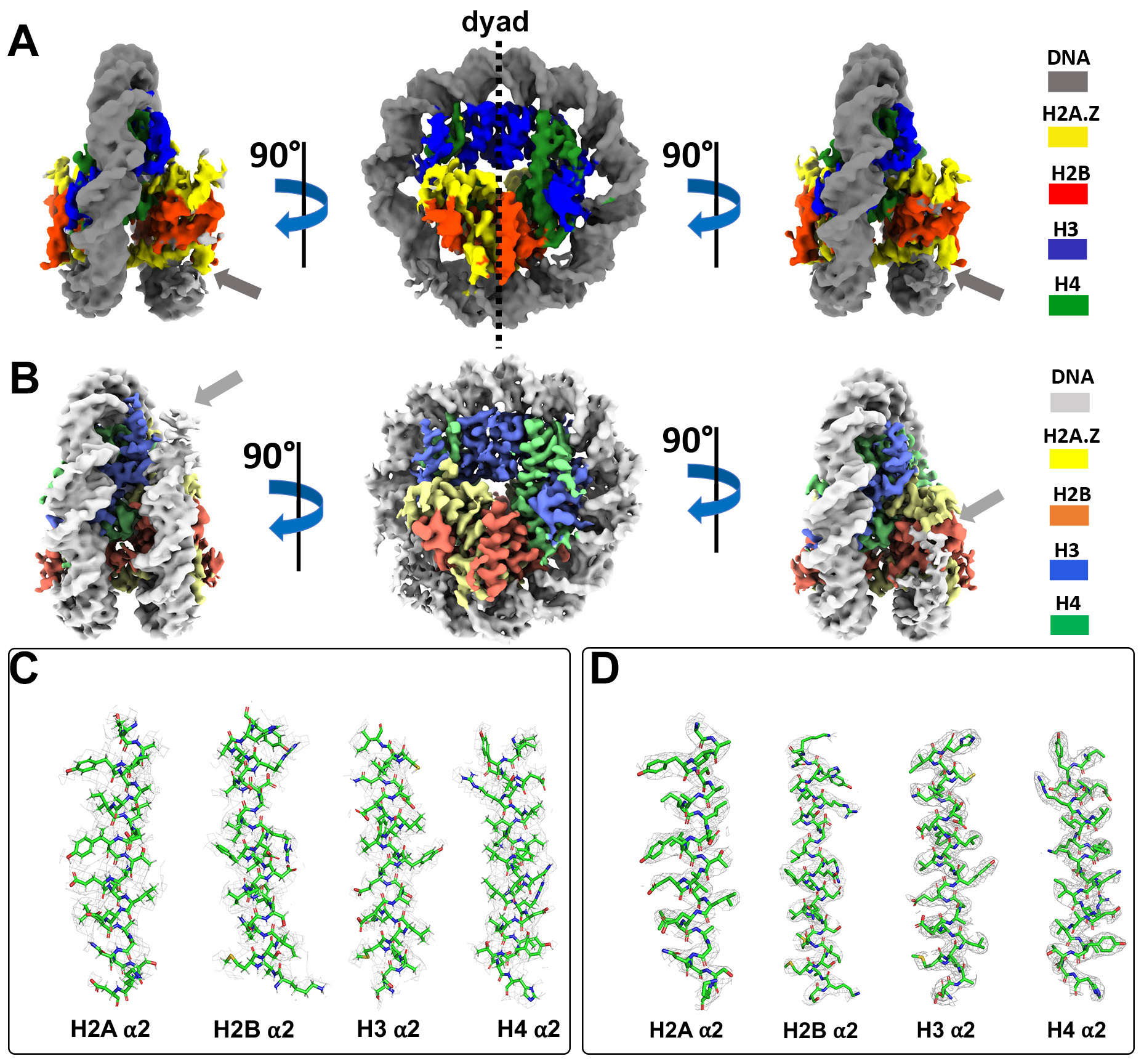

**Extended Data Fig 6. Structural comparison of the K9me3_H2A.Z nucleosomes**

A. Surface representation of the density map (map3 in Extended Data Fig 3C) of the nucleosome in the HP1⍺- H2A.Z complex. Histones and DNA are colored according to the label on the right.
B. Surface representation of the density map of the free K9me3-H2A.Z nucleosome (map shown in Extended Data Fig 5). Histones and DNA are colored according to the label on the right.
C. Close-up view of histone ⍺2 helices in the nucleosome model generated using the map shown in (A).
D. Close-up view of histone ⍺2 helices in the nucleosome model generated using the map shown in (B)

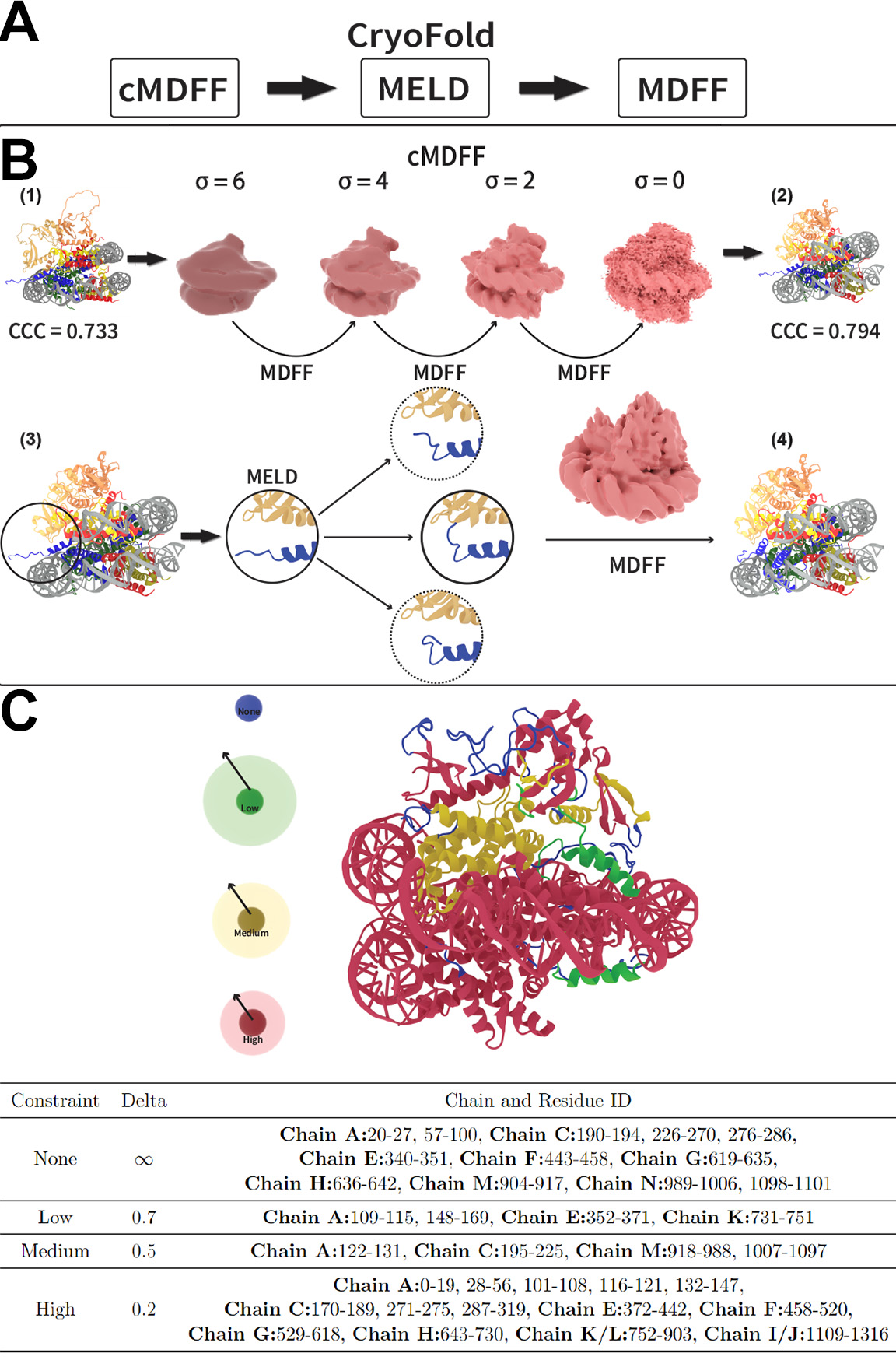

**Extended Data Fig 7. Model building and simulation workflow**

A. Simulation procedure based on the CryoFold algorithm.
B. The initial model (1) with a CCC value of 0.733 underwent a sequential fitting process involving iterative MDFF simulations and a range of smoothed density maps (shown with respective gaussian width values). The resulting model (2) has a CCC value of 0.794. The structure (3) obtained after the cMDFF simulation procedure served as the initial input for MELD simulations which were aimed at refining flexible regions that did not fit the density well. The middle portion highlights the advantage of MELD in that multiple unique conformations of the flexible regions were observed. A final round of MDFF refinement was carried out to obtain the final model (4).
C. **Constraint overview throughout MELD simulations**. The Cartesian restraints (color coded from high to none) were applied to different regions of the model during MELD simulations. The table outlines the extent of the constraints applied to different chains of the model. The delta values (nm) set the threshold of the restraining energy and the allowed displacement range of the alpha carbons. Chain A and C is HP1𝒶. Chain E and K is H3. Chain F and L is H4. Chain G and M is H2A.Z. Chain H and N is H2B.

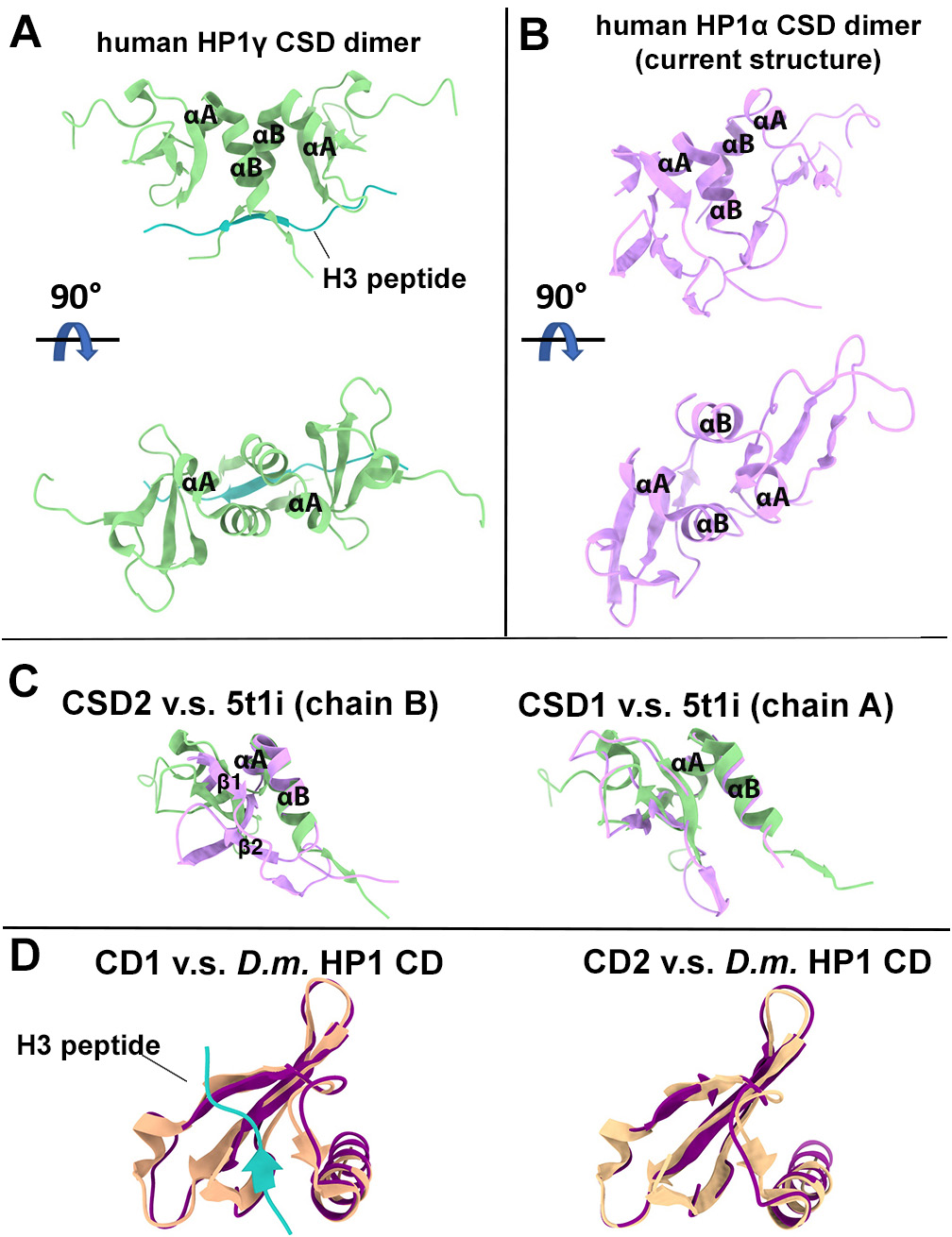

**Extended Data Fig 8 Comparison of the structures of HP1𝒶 domains**

A. Crystal structure of human HP1𝛾 CSD dimer (PDB 5t1i) in complex with an H3 peptide (light blue) in two different views. The four central helices mediating dimerization are indicated.
B. Model of the human HP1𝒶 CSD dimer from the current study in two different view. The four central helices mediating dimerization are indicated.
C. Comparison of CSD2 (left) and CSD1 (right) from the current study (purple) to chain B and chain A of the CSD dimer in the crystal structure (green, PDB 5t1i), respectively.
D. The left panel is the overlay of the crystal structure of *D.m.* HP1 CD (orange, PDB ID 1kne) onto CD1 from the current study (dark purple). Trimethylated H3 tail peptide in the crystal structure is in light blue. The right panel is overlay of the same crystal structure (without the H3 tail peptide) onto CD2 from the current study (dark purple).
